# Comparison of whole-genome bisulfite sequencing library preparation strategies identifies sources of biases affecting DNA methylation data

**DOI:** 10.1101/165449

**Authors:** Nelly Olova, Felix Krueger, Simon Andrews, David Oxley, Rebecca V. Berrens, Miguel R. Branco, Wolf Reik

## Abstract

**Background:** Whole-genome bisulfite sequencing (WGBS) is becoming an increasingly accessible technique, used widely for both fundamental and disease-oriented research. Library preparation methods benefit from a variety of available kits, polymerases and bisulfite conversion protocols. Although some steps in the procedure, such as PCR amplification, are known to introduce biases, a systematic evaluation of biases in WGBS strategies is missing.

**Results:** We perform a comparative analysis of several commonly used pre-and post-bisulfite WGBS library preparation protocols for their performance and quality of sequencing outputs. Our results show that bisulfite conversion per se is the main trigger of pronounced sequencing biases, and PCR amplification builds on these underlying artefacts. The majority of standard library preparation methods yield a significantly biased sequence output and overestimate global methylation. Importantly, both absolute and relative methylation levels at specific genomic regions vary substantially between methods, with clear implications for DNA methylation studies.

**Conclusions:** We show that amplification-free library preparation is the least biased approach for WGBS. In protocols with amplification, the choice of BS conversion protocol or polymerase can significantly minimize artefacts. To aid with the quality assessment of existing WGBS datasets, we have integrated a bias diagnostic tool in the Bismark package and offer several approaches for consideration during the preparation and analysis of WGBS datasets.

## Background

Methylation of DNA at the 5^th^ position in cytosine (5mC) is a stable epigenetic modification found in many living organisms, from bacteria to higher eukaryotes. It is known to play a role in the regulation of transcriptional activity during embryonic development, in processes such as genomic imprinting, transposon silencing, X-chromosome inactivation and during the differentiation of pluripotent cells.

Since its first use in 1992 [1], bisulfite (BS) sequencing of DNA has become the gold standard for analysis of DNA methylation. BS treatment of DNA leads to the conversion of unmodified cytosines to uracil whilst maintaining 5mC unchanged, which, after PCR and sequencing, can be mapped at single base resolution [2,3]. More recently, BS treatment has been coupled with next generation sequencing (NGS) to yield reduced representation (RRBS) or whole genome (WGBS) data on the global genomic distribution of 5mC [4]. As NGS costs decrease, the WGBS approach becomes increasingly accessible for both fundamental and clinical research. However, the ever-increasing diversity of WGBS library preparation kits, protocols and their variations demands a thorough examination of their outputs and performance, to inform the choice of users from both specialist and non-specialist fields, academia and industry. At present, there is a wealth of publically available WGBS datasets, generated in multiple different ways, and it is commonly assumed that they are equally comparable. We set out to investigate how the different steps of current library preparation protocols affect the final sequence output and, ultimately, the quantitation and interpretation of methylation data.

Biases and artefacts from BS sequencing have been well studied outside the NGS context. These encompass biases associated with cloning and PCR, such as primer selectivity and design, polymerase sequence preferences and errors, and template switch (strand recombination) [3,5,6]. In addition, sources of false positive and false negative signals have been well characterised, i.e. the incomplete cytosine conversion by sodium bisulfite and over-conversion of 5mC, found to be affected by factors like DNA quality, quantity and purification procedures, BS incubation length and temperatures, strand reannealing, polymerase, sequencing errors as well as conversion resistant sequences [2,3,5–8]. Different solutions to these biases and artefacts have been proposed, which improved quantitation of DNA methylation at specific loci by PCR and cloning-based methods [2,3,5,7,9–15]. Only some of these considerations, however, remain relevant for NGS-based approaches (e.g. the improvements of BS conversion conditions) and a systematic investigation of major sources of biases in WGBS protocols has not yet been performed. PCR amplification bias has received significant attention in classical (non-BS) whole-genome sequencing [16–22], however, it has been less studied in BS-based whole-genome sequencing [23] and additional sources of bias, which affect both sequencing coverage and methylation quantitation, have not been investigated.

Here we compare several WGBS library preparation protocols by analysing how their sequence coverage and methylation outputs are affected by: 1) BS-induced DNA degradation, 2) PCR amplification, 3) DNA modifications, and 4) incomplete BS conversion. We find that the BS conversion step is the main trigger of biases, due to a selective and context-specific DNA degradation [24] and incomplete conversion efficiency, while subsequent PCR cycles primarily build on the effect of an already biased sequence composition. We discuss mechanisms to avoid, predict or quantitate biases and artefacts in future or for already available WGBS datasets.

## Results

### Study setup: WGBS library preparation steps and strategies

It is well documented that BS conversion causes DNA fragmentation (also known as degradation) of up to 90% of the DNA input [2,7,8,14,24]. In order to assess biases arising from BS-induced DNA degradation, we first tested five bisulfite conversion protocols directly on synthetic and genomic DNA without sequencing. Building on previous work [2,5,7,14], we chose kits from different manufacturers that vary in two key aspects: 1) DNA denaturation, which can be heat-or alkaline-based, 2) BS treatment temperature, which can be high (65-70°C) or low (50-55°C) and typically associated with different incubation times (Table 1). Additionally, we also tested a protocol (‘Am-BS’) that uses high concentration (9 M) of ammonium bisulfite (in contrast with 3-4 M sodium bisulfite used in other protocols) [25].

**Table 1.** BS conversion protocols and parameters.

| Method | Denaturation T°C | Conversion T°C | Incubation Time |
| --- | --- | --- | --- |
| 'Heat 1' | High heat (99°C) | 65°C | 90 minutes<sup>a</sup> |
| 'Heat 2' | High heat (99°C) | 55°C | 10 hours<sup>b</sup> |
| 'Alkaline 1' | Low heat (37°C) | 65°C | 90 minutes<sup>a</sup> |
| 'Alkaline 2' | Low heat (37°C) | 50°C | 12-16 hours<sup>c</sup> |
| 'Am-BS' | Low heat (37°C) | 70°C | 30 minutes<sup>d</sup> |

<sup>a</sup> Imprint DNA Modification Kit (Sigma-Aldrich), 1-step protocol for 'Heat 1' and 2-step protocol for 'Alkaline 1'
<sup>b</sup> EpiTect Bisulfite kit (Qiagen), FFPE protocol and doubled incubation time (see Materials and Methods)
<sup>c</sup> EZ DNA Methylation Kit (Zymo Research)
<sup>d</sup> protocol conditions as described in Hayatsu et al. [25]

We then coupled the above BS conversion protocols to two strategies for the generation of WGBS libraries: 1) pre-BS, which adds sequencing adaptors by ligation before BS conversion [26,27], and 2) post-BS, which adds adaptors by random priming after BS conversion [28]. In total, we tested seven different combinations of BS conversion and library preparation protocols (Table 2). The pre-BS approach involves two DNA fragmentation steps (DNA sonication before library preparation and subsequent BS-induced degradation), and thus requires larger amounts of DNA input (commonly 0.5-5μg). Post-BS approaches overcome this shortcoming, where BS treatment precedes the adaptor tagging and serves to both convert and fragment the DNA, thus utilising only one fragmentation step. This strategy has led to significant reduction in DNA loss and allowed the successful generation of amplification-free WGBS libraries from as little as 400 oocytes [29,30]. Moreover, adding PCR amplification to the original amplification-free post-BS technique allowed sequencing of even lower cell numbers (100-200) and single cells [31–33]. Here we tested the original amplification-free method Post-Bisulfite Adaptor Tagging (PBAT) [28], the PBAT modification with amplification (‘ampPBAT’) [31,32,34], and the commercially available EpiGnome (currently TruSeq) post-BS kit [35,36]. To dissect polymerase differences, we have also included a pre-BS approach performed with the low-bias KAPA HiFi Uracil+ [23] to compare with the most commonly used Pfu Turbo Cx polymerase (Table 2).

**Table 2.**
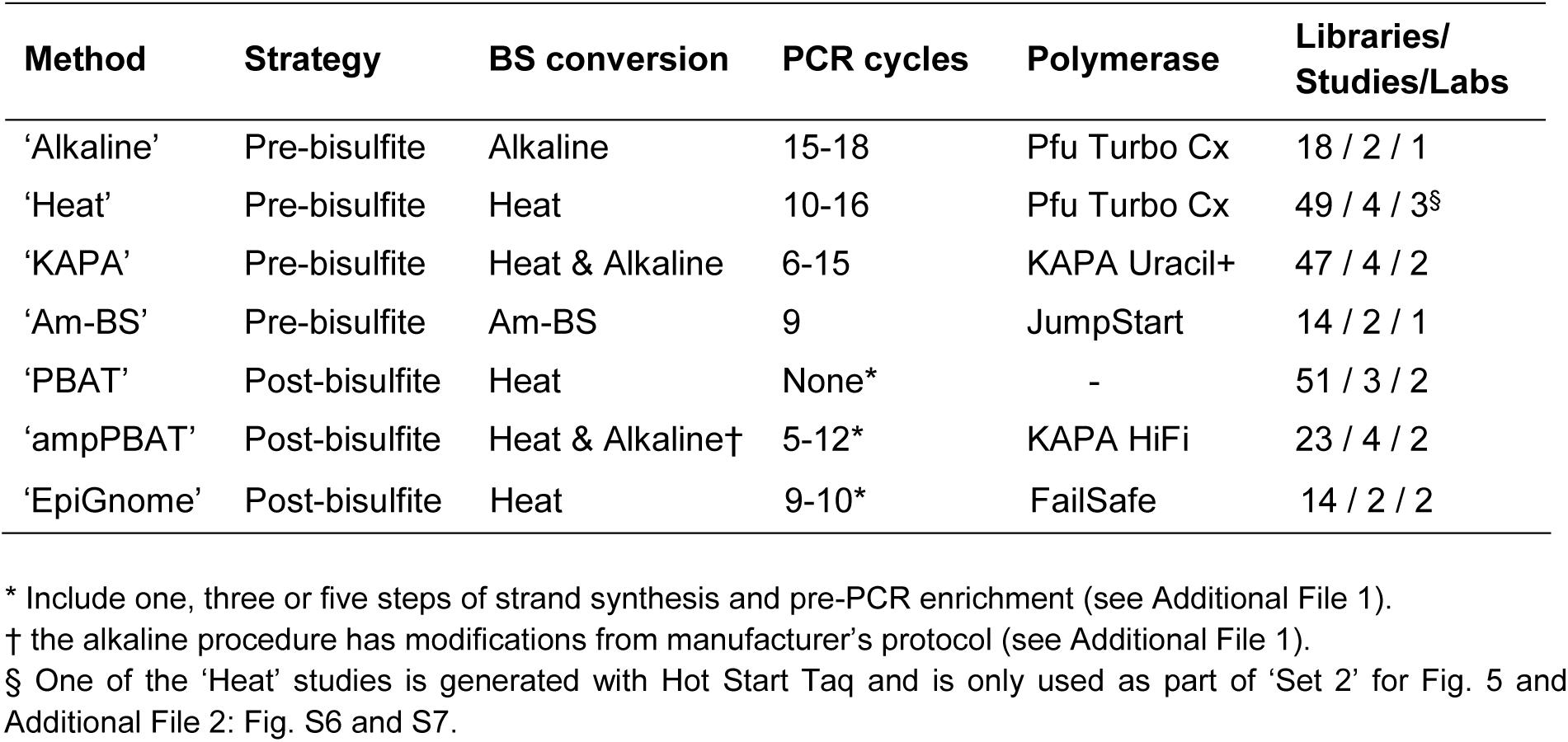
Library preparation parameters of WGBS strategies compared it this study.

In order to increase relevance, comparability and robustness of our study, we have performed a retrospective cross-study and cross-species analysis combining datasets generated by our lab as well as other labs (225 libraries in total including non-BS control; see Table 2 and Additional file 1) [28– 34,36–44]. To capture method differences over batch differences, each method is represented by at least two studies sourced by two different laboratories, where possible (Table 2). This ensures that the trends described herein are observed across data generated by a wide scientific community and not inherent to a single lab’s results.

### Effect of BS-induced DNA degradation

DNA degradation is a well-known concomitant effect of BS conversion, which has made challenging its usability for low cell numbers, but has never been reported as a factor creating sequence biases. BS-induced fragmentation was initially attributed to loss of purines [1,7], but was later shown to result from random base loss at unmethylated cytidines, which causes backbone breakage upon exposure to heat and alkali [24]. Such cytosine-specific effect could lead to two possible biases: 1) depletion of cytosine-rich DNA from the total sequence pool, resulting in a skewed representation of genomic sequences, and 2) depletion of unmethylated fragments, leading to an overestimation of the absolute 5mC values. To test these possibilities, we BS treated synthetic DNA fragments of low (15%, ‘C-poor’) or high (30%, ‘C-rich’) cytosine content (see sequences in Additional file 2: Table S1). Strikingly, the recovery of the C-poor fragment was 2-fold higher than that of the C-rich fragment when using the ‘Heat’ BS treatment (Fig. 1a). The milder ‘Alkaline’ denaturation showed higher recovery and reduced bias across cytosine contents (1.3-fold difference), whereas the ‘Am-BS’ protocol showed no significant difference between C-contents, despite its relatively low recovery (Fig. 1a). Reducing BS incubation temperature from 65°C to 50-55°C at the expense of longer incubation (Table 1) did not produce a difference in yields (Additional file 2: Fig. S1a). Results from ‘Heat’ 1 and 2 or ‘Alkaline’ 1 and 2 pairs have therefore been pooled as ‘Heat’ and ‘Alkaline’ in subsequent analyses, unless otherwise stated. These results suggest that BS conversion conditions could have an impact in genomic coverage in WGBS.

**Figure 1.**
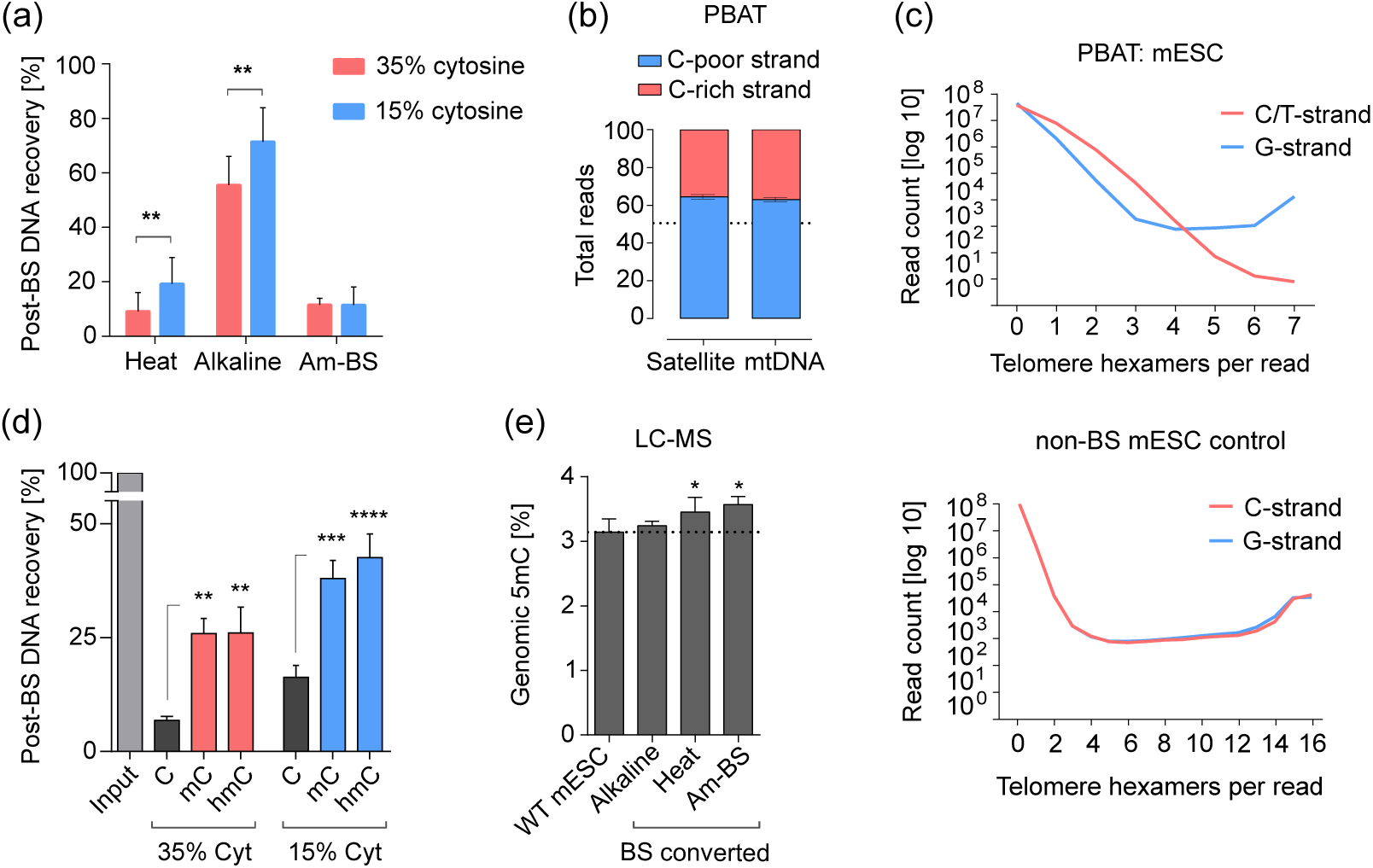
Biased degradation of unmethylated C-rich DNA after bisulfite treatment. (**a**) Post-bisulfite recovery of C-rich and C-poor DNA fragments treated with different BS conversion protocols. Fragment sequences originate from the M13 phage sequence (Additional file 2: Table S1). Statistical analysis was performed with a two-way ANOVA, p = 0.0034 for cytosine content (‘Heat’ and ‘Alkaline’ only) and p < 0.0001 for the method. (**b**) Asymmetric C-rich (23-24% C) and C-poor (12-14% C) strand representationin the mouse major satellite repeat and mtDNA in amplification-free PBAT datasets. The total read count per strand is represented as a proportion of100%. (**c**) Telomere repeat count per read in amplification-free PBAT and non-BS converted NGS control. To assess the fragmentation rate of the C-strand with increase of tandem count and cytosine content, reads containing G-strand tandems ([TTAGGG] _n_) were quantitated separately from unconverted and BS converted (in PBAT) C-strand tandems ([CCCTAA] _n_ and [TTTTAA] _n_, respectively). Each plot represents a single dataset. C-strand reads containing less than four to six tandems are genuine [TTTTAA] _n_repeats of non-telomere origin (results not shown, see Methods). (**d**) Post-bisulfite recovery of unmethylated, fully methylated and hydroxylated C-rich and C-poor DNA fragments treated with different BS conversion protocols. One-way ANOVA was performed with Tukey’s multiple comparisons test. (**e**) LC/MS measurement of total genomic 5mC levels in mouse ESC gDNA before (‘WT mESC’) and after treatment with different BS-conversion protocols (‘Alkaline, ‘Heat’, Am-BS’). 5mC is represented as a percentage of all cytosines; the converted cytosines were measured as uracils in the BS converted samples. Individual two-tailed paired t-tests were performed within matched sample-control pairs. Details on number of WGBS datasets used for each analysis are presented in Additional File 1. *p<0.05, **p<0.01, ***p<0.001, ****p<0.0001; error bars represent s.d.

To test whether DNA degradation leads to uneven sequence coverage in WGBS data, we sought genomic regions where the relative strand coverage could be affected by the depletion of cytosines. Both the major (pericentric) satellite repeat and mitochondrial DNA (mtDNA) display substantial differences in cytosine content between their lower and upper strands (Additional file 2: Table S2). To exclude interference of PCR bias, we only analysed amplification-free PBAT datasets, which employ heat-based DNA denaturation (Table 2 and Additional File 1). Both mouse major satellites and mtDNA showed significantly higher coverage of their C-poor (12-14% cytosine) in comparison to their C-rich (23-24% cytosine) strand (Fig. 1b). We also examined the telomere repeat, comprised of 50% cytosines on one strand ([CCCTAA]_n_, ‘C-strand’) and none on the other ([TTAGGG]_n_, ‘G-strand’). BS sequencing reads from high copy number tandem telomere repeats showed up to 1000-fold higher coverage of the G-strand compared to the C-strand (Fig. 1c and Additional file 2: Fig. S1b). Notably, whilst BS sequencing cannot distinguish BS-converted CCCTAA repeats from genomic TTTTAA repeats, the abundance of the latter in the lower copy numbers (< 4-5 repeats in figure) cannot explain the observed bias in the high copy numbers. These results confirm that the WGBS output of unmethylated C-rich sequences is affected by BS-induced degradation.

To test the effect of cytosine modifications on DNA fragmentation, we generated 5mC-and 5hmC-modified C-poor and C-rich fragments. Both modifications yielded a ∼4-fold increase in recovery for the C-rich sequence and more than two-fold for the C-poor sequence with the ‘Heat’ BS-conversion protocol (Fig. 1d). A weaker protective effect on the C-rich fragment was observed for the ‘Alkaline’ conversion protocol, whereas the ‘Am-BS’ protocol showed 3-4 fold increase in recovery for both fragments (Additional file 2: Fig. S2a). This indicates that cytosine modifications have a protective effect against BS-induced DNA degradation, especially in C-rich sequences. Finally, analysis of BS converted DNA from mouse embryonic stem cells (mESC) by liquid chromatography coupled to mass spectrometry (LC/MS) revealed that both highly degrading ‘Heat’ and ‘Am-BS’ protocols cause a direct 5-10% increase in the global estimate of DNA methylation, whilst no such effect was observed for the milder ‘Alkaline’ procedure (Fig. 1e). Differences in DNA clean-up procedures affected overall yields (Additional file 2: Fig. S2b), but were not responsible for the observed differences in the estimation of methylation by LC/MS (Additional file 2: Fig. S2c).

In summary, BS-induced DNA degradation leads to depletion of genomic regions enriched for unmethylated cytosines, which creates a biased sequence representation and directly affects the final estimation of 5mC levels. DNA degradation is strong in harsher BS conversion protocols that utilise high denaturation temperatures (‘Heat’) or high BS molarity (‘Am-BS’).

### Effect of PCR amplification bias

PCR amplification is a notorious source of bias in massively parallel sequencing, known to affect primarily sequences with highly skewed base composition on the extreme ends of GC content [18]. This has led to technical difficulties, which have partially been resolved by new amplification-free approaches [22], PCR buffer additives [16,18–20], better temperature control over PCR steps and cycle ramp rates [16,21], and extensive screens for low bias polymerases [16–20,23]. The mammalian BS converted genome has ∼80% AT content and ∼20% G content, which makes it a real challenge for polymerases. Although the commonly used Pfu Turbo Cx polymerase is not the worst performer among its counterparts [17], the KAPA HiFi family of polymerases used together with the PCR additive TMAC (tetramethylammonium chloride) have shown the best tolerance for AT-rich regions, albeit at the expense of higher error rates [18–20,23]. A more recent study suggested that the bias resulting from PCR in NGS stems primarily from factors such as stochasticity of amplification of low-copy sequences and polymerase errors [21]. These factors seem especially relevant for BS converted DNA, given the high degradation of input material and the high reported rate of polymerase sequencing errors in high AT content DNA.

To evaluate the effect of PCR biases on sequence representation, we quantitated the dinucleotide coverage in all datasets from a cross-laboratory panel of WGBS methods (Table 2 and Additional file 1) against the expected genomic value (Additional file 3) and compared to a non-converted control. All methods showed a highly significant dinucleotide coverage bias relative to control (Fig. 2a and Additional file 2: Fig. S3a). The observed biases were largely consistent across multiple libraries from different laboratories, although the extent of these biases varied somewhat between studies (Additional File 2: Fig. S3a). There was a clear enrichment for G-containing dinucleotides and depletion of AT-rich sequences in all methods, with the exception of ‘KAPA’, which showed a balanced G content, an unexpected depletion of C content and enrichment of AT content (Fig. 2a and Additional file 2: Fig. S3a). These features of the ‘KAPA’ profile were not affected by the BS conversion protocol used and the bias did not decrease in libraries generated with fewer PCR cycles (Additional file 2: Fig. S3a). A similar result was observed for the post-BS ampPBAT method (Additional file 2: Fig. S3a). Interestingly, the amplification-free PBAT also showed a slight G-bias, possibly due to its DNA synthesis and pre-PCR enrichment steps (Fig. 2a, right panel). However, unlike other methods, amplification-free PBAT did not display any significant deviation from control with respect to CG-dinucleotide coverage, where most methylation occurs, suggesting that the amplification-free post-BS approach has the lowest methylation biasing (Fig. 2b, left panel). CG coverage varied highly overall, and for pre-BS methods it seemed more affected by polymerase than by BS conversion protocol, while for post-BS ‘ampPBAT’ the Heat BS treatment produced significantly over-represented CG over ‘ampPBAT’ Alkaline BS converted libraries. Such an effect was not observed for the closest structurally CA dinucleotide (Fig. 2b, right panel).

**Figure 2.**
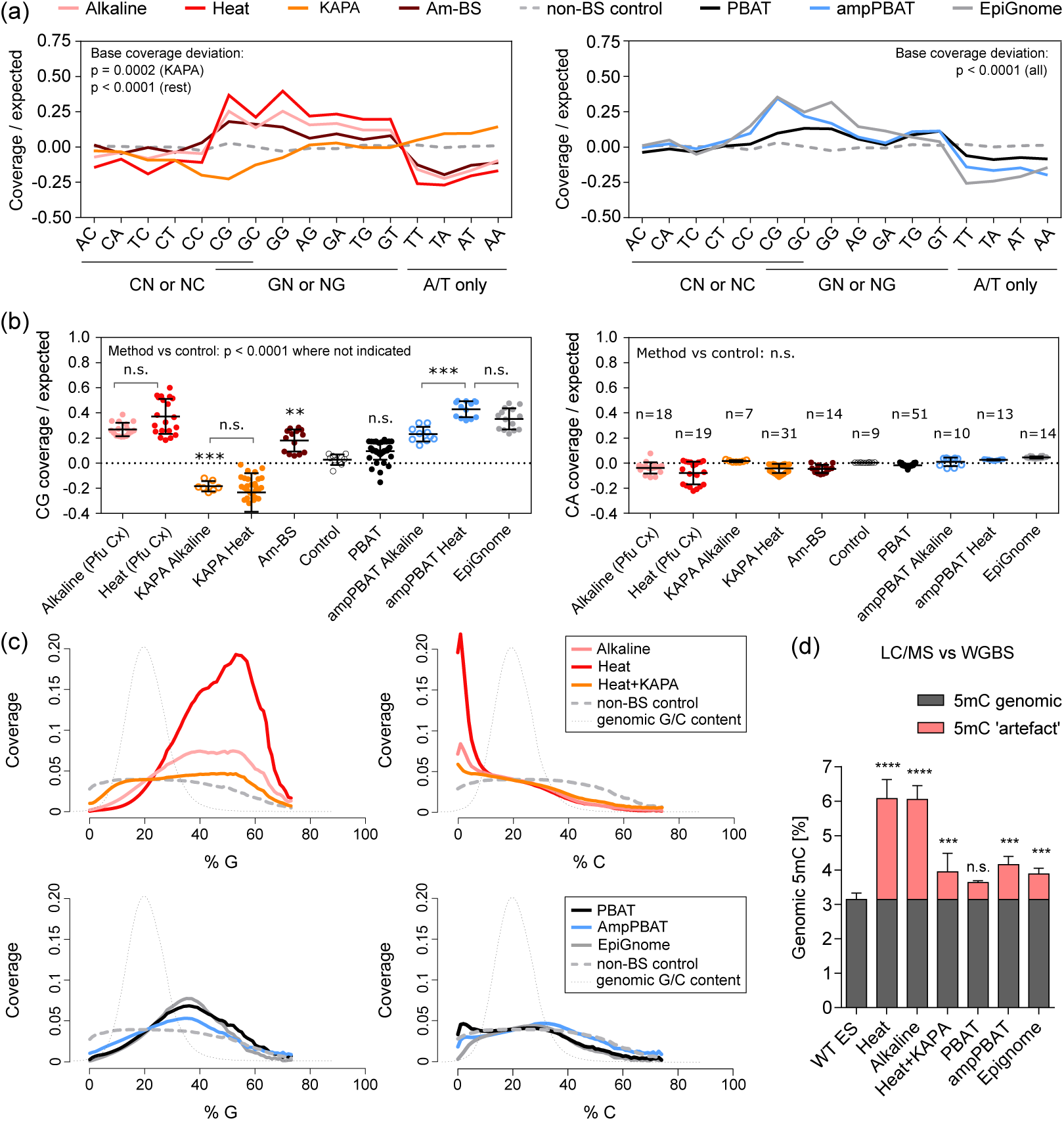
Effect of PCR and polymerase bias on sequence coverage and methylation estimates. (**a**) Coverage of dinucleotides in WGBS datasets generated with our pre-bisulfite (left panel) and three post-bisulfite (right panel) library preparation protocols. Averaged coverage values per method are expressed as log2 difference from the genomic expected and compared to a non-BS-treated control. For clarity, the dinucleotides are underlined as derived from C, G, or A /Tonly. Statistical analysis of overall dinucleotide (base) coverage was performed on the average absolute deviations from control with one sample wo-tailed t-tests followed by Bonferroni correction. Individual libraries/sequencing runs and studies per method are presented in Additional file 2: Fig. S3a;details on study, laboratory and species can be foundi n Table 2 and Additional File 1. (**b**) CG dinucleotide coverage over expected for each individual library per method (left panel) and unbiased CA dinucleotide coverage for comparison (right panel). The main method groups are additionally split into subgroups of Heat and Alkaline for KAPA and ampPBAT, as shown also in Additional file 2: Fig. S3a. The number of libraries per method is presented in the right panel. ‘Pfu Cx’ in brackets next to the main ‘Alkaline’ and ‘Heat’ methods stands for the Pfu Turbo Cx polymerase, which they are generated with (all method details are provided in Table 2). Statistical analysis on CG coverage was performed with one-way ANOVA with Bonferroni correction. (**c**) Read coverage dependence on the G/C composition in 100 bp tiles. Cytosine content of reads was calculated from the corresponding genomic sequence and not the actual read sequence, where unmethylated cytosines appear as thymines. The tile distribution per G/C content is plotted in the background along the x-axis for ref erence. Am-BS was omitted from this analysis due to unavailability of same species datasets. (**d**) Global methylation levels of mouse ES cells as measured by LC/MS and a panel of WGBS datasets. The LC/MS value is an average for J1 and E14 lines from different passages and studies to account for lineage and tissue culturing variances. The differences between the LC /MS values and WGBS measurements are marked as a methylation ‘artefact’ within each WGBS method. Significance is calculated by one-way ANOVA with Dunnett’s multiple comparisons test on the absolute WGBS values (‘genomic’ + ‘artefact’) against the LC/ MS value; **p<0.01, ***p<0.001, ****p<0.0001; error bars represent s.d.

We next asked in more detail how sequencing depth was affected by the % of G or C. These analyses confirmed that KAPA-amplified libraries displayed the lowest amount of G content bias, in contrast to the drastic G-enrichment of Pfu Turbo Cx’s ‘Heat’ and ‘Alkaline’ pre-BS datasets (Fig. 2c upper panels). The post-BS methods also showed a more balanced G coverage, performing similarly regardless of amplification (Fig. 2c lower panels). With respect to C coverage the post-BS methods outperformed the pre-BS group, where both ‘KAPA’ and Pfu Turbo Cx’s ‘Heat’ and ‘Alkaline’ datasets under-represent the C-high (> 25% C) and over-represent the C-low (< 15% C) sequences. This suggests that the BS-degradation bias, characterised by depletion of C-rich sequences, does not affect post-BS approaches to the extent that it affects pre-BS protocols.

Next, we investigated how polymerase bias affected regions with known GC-skew between strands. For both the satellite repeat and mtDNA, PCR amplification exacerbated the strand bias arising from DNA degradation observed in ‘PBAT’ in Fig. 1b (Additional file 2: Fig. S3b), demonstrating a combined effect of the PCR and degradation biases (lowest for the ‘KAPA’ datasets). Similarly, the C-rich strand of the telomere repeat was hardly detectable in the raw reads of most datasets (Additional file 2: Fig. S3c).

Finally, to assess the effect of amplification on quantitation of 5mC, we quantitated total 5mC levels in mESC within our panel of WGBS datasets (except for ‘Am-BS’, for which mESC data was not available). The levels in both ‘Heat’ and ‘Alkaline’ datasets doubled the LC/MS-measured 5mC value, whilst the overestimation in the ‘KAPA’ and post-BS datasets was less pronounced (Fig. 2d). Notably, 5mC levels in amplification-free PBAT were not significantly different from those measured in genomic DNA (Fig. 2d) and were comparable to those detected after BS conversion alone (Fig. 1e).

In summary, all WGBS approaches show significant sequence coverage deviations in comparison to conventional WGS. Amplification-free PBAT, however, shows better representation of C-rich and C-low sequences, resulting in a less pronounced overestimation of global methylation values.

### Effect of DNA modifications

To further evaluate the genome-wide influence of DNA modifications on WGBS biases, we compared gDNA from DNMT-Triple Knockout (TKO) mESCs, which lack DNA methylation, to *in vitro* methylated TKO gDNA (‘meTKO’) using the M.CviPI methylase. We sequenced TKO and meTKO WGBS libraries generated using the ‘Heat’ protocol [45], which was strongly affected by depletion of C-containing sequences in our previous analyses (Fig. 2a and 2c). The meTKO library displayed ∼20% methylation across all cytosine contexts (Additional file 2: Fig. S4a) and this translated into a ∼15% higher sequencing coverage at all cytosine-containing dinucleotides when compared to the TKO library (Fig. 3a). This shows that DNA methylation affects sequence coverage, as suggested by our experiments with modified DNA fragments (Fig. 1d). Interestingly, differences in total 5mC levels between meTKO and WT mESC DNA (15% versus 3%, by LC/MS) led to differences in the overestimation of 5mC by WGBS (40% versus 100% overestimation; Fig. 3b and 2d), presumably because meTKO DNA has fewer unmethylated cytosines available for BS-induced degradation. The accuracy of local 5mC measurements by ‘Heat’ WGBS therefore depends on the extent of methylation at each locus. We also found that DNA methylation affected read coverage depending on the % of C or G content (Fig. 3c). Notably, the even distribution of *in vitro* deposited 5mC was associated with a more even coverage across C/G content when compared to TKO DNA (Fig. 3c), suggesting that a substantial amount of the coverage bias in WT mESCs (Fig. 2c) is driven by differences in the genomic distribution of 5mC. An increase in coverage, albeit weak, was also observed for the satellite repeat and mtDNA C-rich strands (Additional file 2: Fig. S4b).

**Figure 3.**
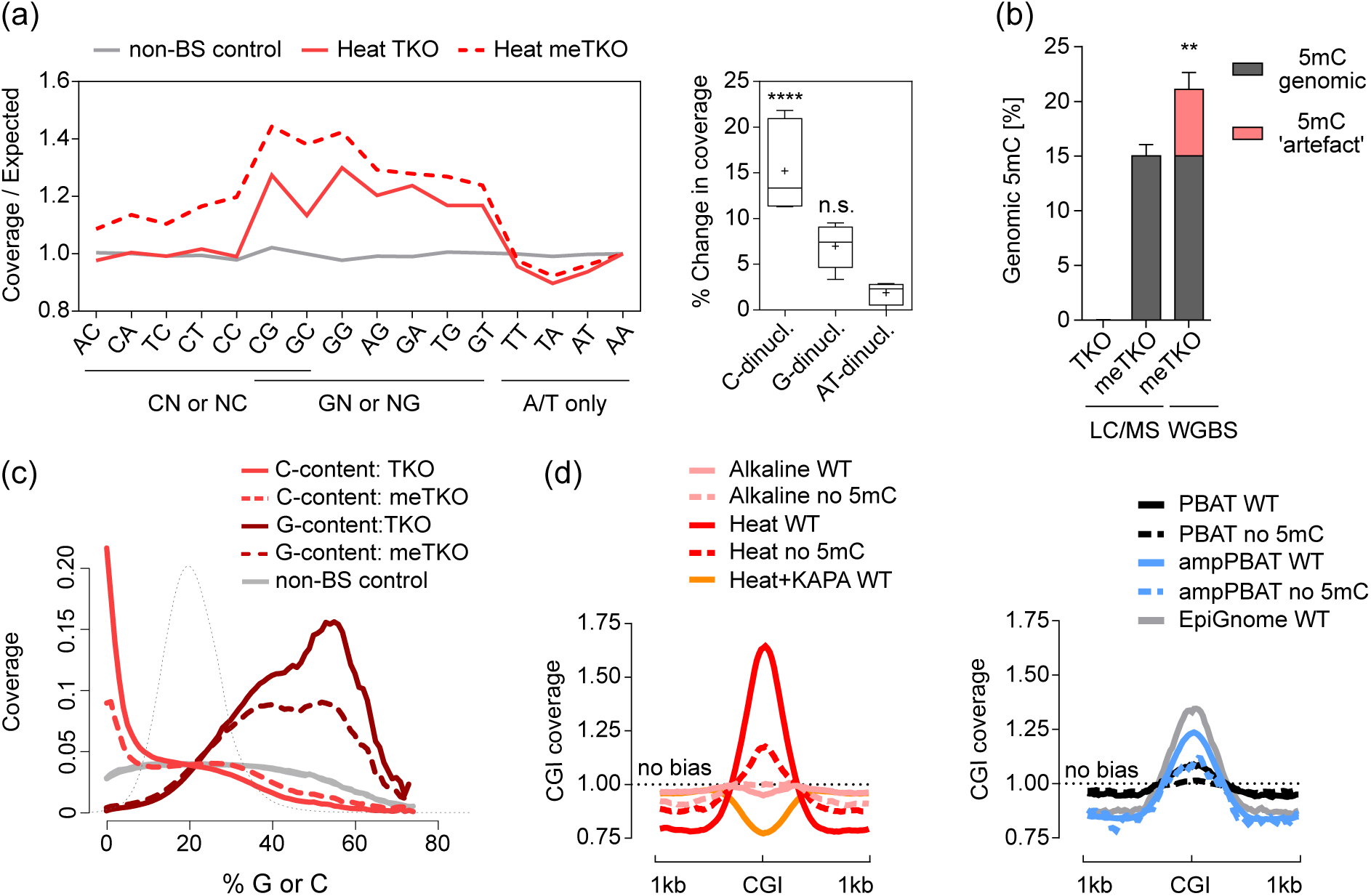
Effect of DNA methylation status on the degradation and amplification biases. (**a**) Coverage of dinucleotides in WGBS datasets from unmethylated and *in vitro* M.CviPI-methylated TKO DNA prepared with the ‘Heat’ BS-seq protocol. For direct comparison, the increase in coverage is expressed as fold difference from the genomic average and normalised to the AA dinucleotide. The dinucleotides are grouped as derived from C, G or A/T only and presented in the box-plot panel (right) as total % increase in coverage; crosses mark mean values and error bars represent minimum and maximum values. Statistical analysis was performed with one-way ANOVA with Dunnett’s multiple comparisons test against the AT-only dinucleotides; ****p<0.0001. (**b**) Global methylation levels of the *in vitro* M.CviPI-methylated TKO DNA. The difference between the LC/MS and WGBS values is marked as a methylation ‘artefact’ as in Fig. 2. Significance between the two values was assessed with a two-tailed unpaired t-test, p=0.0011, error bars represent s.d. (**c**) Read coverage dependence on the G/C composition before and after *in vitro* M.CviPI methylation. Cytosine content of reads was calculated over 100 bp tiles and matched to the corresponding genomic sequence and not the actual read sequence, where unmethylated cytosines are converted to thymines. Dotted black line represents the tile count distribution in G/C content. (**d**) Coverage of CGIs in WGBS datasets of mouse WT ESCs (i.e. with similar level of methylation) compared to unmethylated DNA generated with the same library preparation protocols. Heat+KAPA and EpiGnome protocols are included only as WT values for reference, due to unavailability of corresponding unmethylated DNA datasets. Values are expressed as fold difference from a coverage ‘no bias’ line.

These results suggest that highly methylated sequences could be over-represented in WT genomes. To explore how methylation status affected GC-rich regions of interest, we compared averaged CGI coverage in our panel of mESCs WGBS datasets. Low levels of 5mC seen at CGIs have the potential to cause a coverage bias in comparison to entirely unmethylated genomes; therefore we also included unmethylated samples for all methods apart from ‘KAPA’ and ‘EpiGnome’(which were not available). The ‘Heat’ pre-BS method showed highest coverage bias over WT mESC CGIs, which decreased significantly in the unmethylated sample (Fig. 3d). The post-BS methods showed low (‘PBAT’) to moderate (‘ampPBAT’ and ‘EpiGnome’) coverage bias, which also decreased in the unmethylated samples (Fig. 3d, right panel). No methylation-dependent coverage bias was detected with the ‘Alkaline’ protocol, and an under-representation of CGIs was observed for the AT-biasing ‘KAPA’ datasets. These results confirm that particular C/G-rich sequences will be over-or under-represented in WGBS datasets, depending on their methylation status and the chosen library preparation strategy.

### Effect of incomplete BS conversion

It is known that different BS treatment conditions have variable efficiency in converting cytosine to uracil [2,5,14]. Given the variable preference of polymerases towards cytosine and the depletion of successfully converted cytosines through BS-induced degradation, we asked whether unconverted cytosine artefacts could contribute to the observed overestimated methylation values in WGBS datasets (Fig. 2d). To measure conversion efficiency of the BS conversion protocols in Table 1, we BS-treated mESC gDNA and quantified the amount of unconverted cytosines by LC/MS (LC/MS can distinguish unconverted Cs from 5mCs). Our results show that ‘Heat’ denaturation yielded best conversion, whereas ‘Alkaline’ denaturation led to 4-fold higher amounts of unconverted cytosine (Fig. 4a). When added to the total 5mC levels measured by LC/MC (Fig. 4b) it is clear that unconverted cytosines make a substantial contribution to the overestimation of 5mC levels by WGBS (Fig. 2d). This effect will be strongest for the milder ‘Alkaline’ conditions, which was confirmed in our WGBS datasets of the same unmethylated TKO samples prepared with the ‘Heat’ and ‘Alkaline’ methods (Fig. 4c). Strikingly, the % of uncoverted cytosines in the ‘Alkaline’ TKO datasets surpassed the biological value of real 5mC levels in WT mESCs, demonstrating how unconverted cytosines can lead to vastly increased total 5mC estimates.

**Figure 4.**
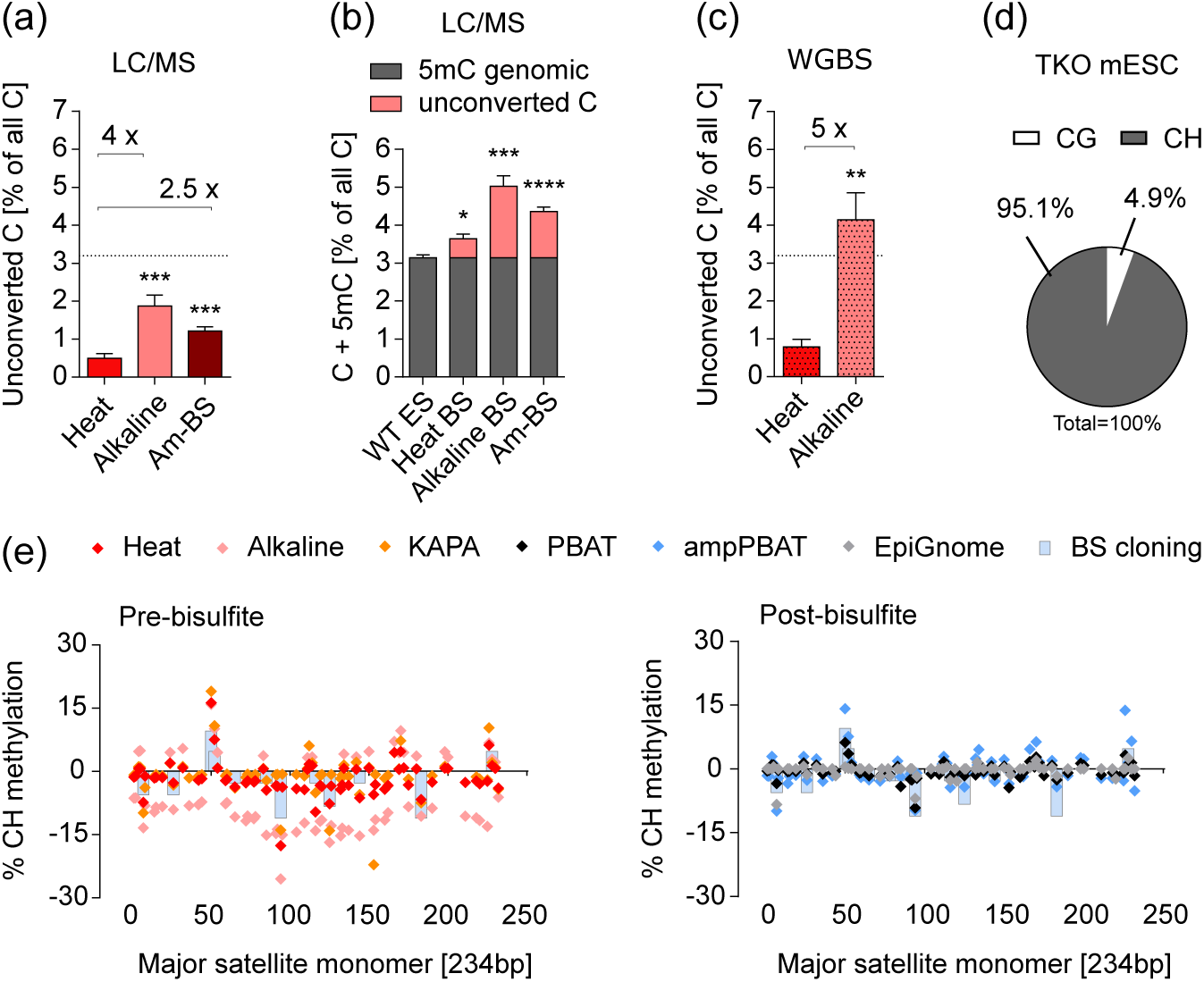
Effect of conversion artefacts on the biases in WGBS. (**a**) Presence of unconverted cytosines as percent of total cytosine content, measured by LC /MS for three different BS-conversion protocols. The three protocols differ by denaturation method (‘Heat’ or ‘Alkaline’) or molarity of bisulfite (4.5M vs 9M for ‘Am-BS’) but not by BS incubation temperature (65-70°C). Averaged fold-differences in quantity are shown above horizontal brackets, and a dotted line shows the usual level of genomic 5mC for reference of scale. For conversion differences between methods with 50°C and 65°C incubation temperatures, see Additional file: Fig. S10a. (**b**) A theoretical sum of 5mC and unconverted C as measured by LC/MS for J1 WT mESCs for three BS conversion protocols. Both 5mC and unconverted C will be interpreted as 5mC after amplification of WGBS libraries, boosting the overall levels of methylation, depending on the BS treatment protocol. (**c**) Absolute quantitation of unconverted cytosines in the unmethylated TKO mESC line, as measured by ‘Heat’ and ‘Alkaline’ BS-seq.(**d**) Context distribution of BS conversion artefacts; the valueis the same for ‘Heat’ and ‘Alkaline’and therefore plotted as an average. (**e**) CH methylation on both strands of the mouse major satellite repeat as measured by pre -and post-bisulfite WGBS methods. 5mC percentage from the BS cloning from Additional File: Fig. S5a is plotted in both panels for reference. Positive y-axis values indicate the top strand and negative – the bottom strand. Statistical analyses in (a), (b), and (c) were performed for matched experimental pairs with unpaired two-tailed t-tests against ‘Heat’ in (a) and (c), and ‘WT ES’ in (b). Error bars in (a), (b) and (c) represent s.e.m.,*p < 0.05, **p<0.01, ***p<0.001, ****p<0.0001.

Conversion artefacts occur mainly in non-CG context (or CH, where H is A, T or C) (Fig. 4d). This is because CH context is over 20-fold more abundant in mammalian genomes than the CG context (Additional file 3). We therefore sought examples of genomic locations that could be particularly affected by this artefact. Mouse major satellites have been shown to have non-CG context methylation through a BS cloning based approach [46]. As targeted BS sequencing relies on primers that select for fully converted fragments [9], these results are more likely to reflect the real non-CG methylation distribution. We therefore produced our own BS cloning data for major satellites to compare against WGBS results (Fig. 4e and Additional file 2: Fig. S5a,b). Analyses of our panel of WT mESC WGBS datasets showed that, unlike what is seen by BS cloning, pre-BS ‘Alkaline’ datasets had pronounced strand asymmetry in the CH methylation levels, which were more abundant on the C-rich (bottom) than the C-poor (top) strand (Fig. 4e, left panel). The ‘Heat’ protocol had a reduced bias, whereas the ‘KAPA’ method did not show strand asymmetry, although certain positions had higher mCH values than those obtained by BS cloning. None of the post-BS methods showed strand asymmetry, even after amplification (Fig. 4e, right panel). These results can be explained by the preferential amplification of poorly converted reads, which are less likely to be degraded, whereas well converted C-rich reads from the bottom strand tend to be degraded, as we have shown. This gives the false perception of asymmetric methylation in such regions. Indeed, asymmetrically methylated regions are commonly reported in BS-seq datasets, especially in CH context [47]. Notably, mCG levels were also asymmetrically elevated, reaching 20% in some instances (Additional file 2: Fig. S5c). These results illustrate a direct link and interplay between sequence-specific BS-induced degradation, conversion errors and the amplification of those artefacts by PCR, leading to higher overall bias in protocols with amplification.

### Effect on quantitation of methylation

To investigate how the different biases affect the final quantitation of methylation in genomic features we analysed two sets of data: 1) Set 1 included the previously used public mESCs WGBS datasets prepared with six different protocols by different teams [29,31,36,39,40,48,49] and 2) Set 2 included a combination of ‘PBAT’ and BS-seq ‘Heat’ datasets from four different biological samples (mESC, blastocyst, oocyte and sperm) prepared by a single team [29].

We have already shown how the different methods in Set 1 performed in estimating global 5mC levels (Fig. 2d) and with respect to CG coverage (Fig. 2a-c). The Set 2 ‘Heat’ protocol uses a different polymerase that also leads to a depletion of C-containing dinucleotides (Additional File 1: Fig. S6a). To test whether this indeed translates into higher local CG methylation, we first looked at the methylation levels of imprinted differentially methylated regions (iDMRs), which are expected to have mCG values close to 50% in mESCs. We found substantial differences in DMR methylation values, ranging from 44% in ‘PBAT’ (which showed a global 5mC value nearest to the LC/MS value in Fig. 2c) to 58% in the EpiGnome protocol (Fig. 5a). ‘PBAT’ consistently yielded the lowest methylation values also across multiple genomic features such as genes, intergenic regions and repeats, whereas the rest of the methods showed comparably higher mCG values for all but the highly methylated IAP repeats (Additional file 2: Fig. S6b). This prompted us to test whether the methylation increase over the ‘PBAT’ values was linear across all 5mC values, for which we did a genome-wide comparison of each method against PBAT. This revealed that the largest discrepancies lay in the middle range of CG methylation values, i.e. the moderately or variably methylated genomic regions (Fig. 5b and Additional file 2: Fig. S6c). The same result was obtained when comparing ‘Heat’ and ‘PBAT’ data for the four biological samples from Set 2, but not for the individual ‘Heat’ replicates, validating that the observed non-linear discrepancies are not due to batch effects or lab-to-lab variability (Additional file 2: Fig. S6d). Importantly, moderately methylated regions are commonly studied targets in biological samples, since they indicate areas of variability and heterogeneity within an epigenetic pool and often include enhancers, promoters and transcription factor binding sites [50]. A closer look into such features in Set 1 confirmed that they display larger methylation differences between PBAT and the other methods when compared to genic and repeat regions (Fig. 5c and Additional file 2: Fig. S7a-c). These results further highlight that some regions are more susceptible to biases than others and the method variability in estimation of 5mC across the genome does not follow a linear fashion.

**Figure 5.**
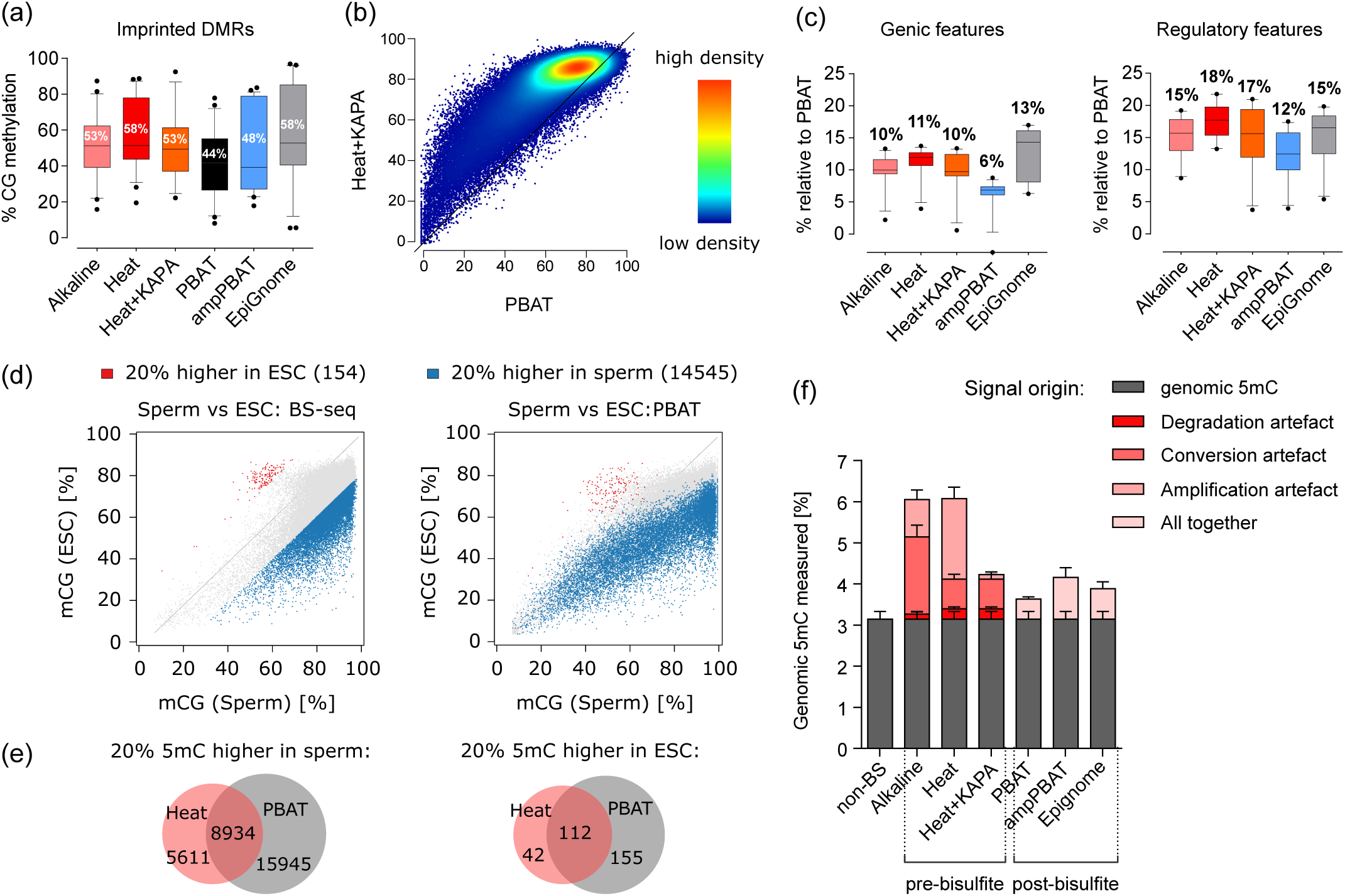
Effect of biases and artefacts on the output of 5mC quantitation. (**a**) Average methylation values of iDMRs. Numbers indicate the mean value, error bars span the 10-90 percentile. (**b**) Genome-wide comparison of absolute methylation levels for the amplification-free PBAT approach and the ‘Heat+KAPA’ BS-seq. (**c**) Differences in the absolute quantitation of genomic and regulatory features between the amplification-free PBAT (at value 0) and amplified WGBS datasets. Numbers indicate the mean value, error bars show the 10-90 percentile. (**d**) Comparison of relative methylation differences between sperm and ESC sequenced with either amplification-free PBAT or ‘Heat’ BS-seq. Each dot represents a probe over 150 consecutive cytosines from the same genomic region in ESC and sperm. The plotted over 20% mCG differences are generated from the BS-seq method (left panel) and visualised with same colour onto the PBAT data (right panel). Averaged values were used for BS-seq (2 x ESC and 5 x sperm replicates) and a single replicate for PBAT. (**e**) Venn diagrams showing how many of the over 20% mCG regions from (d) overlap bet ween BS-seq and PBAT. (**f**) A breakdown of relative contribution of biases for the BS-seq protocols as measured by LC/MS and WGBS. For post-bisulfite pro tocols, the overall combined bias is shown as individual contributions are less trivial to dissect. The non-BS 5mC measurement averages LC/MS measurements for mES lines from different studies and passages to account for culturing and lineage variances. Error bars represent s.d.

Despite the clear differences in absolute methylation values between the WGBS methods, DNA methylation is often studied as a relative change between conditions, treatments and biological samples. To address whether relative changes in methylation differ when analysed with pre-or post-BS WGBS, we analysed methylation differences between BS-seq and PBAT datasets from sperm and ESC from Set 2. We selected regions with more than 20% methylation difference between the two samples in both directions from one of the methods, and compared the positioning of those regions in the other method (Fig. 5d and Additional File 2: Fig. 8a). More than half of the regions identified with the ‘Heat’ method were also identified with ‘PBAT’ but the larger proportion of regions from ‘PBAT’ were not identified with ‘Heat’ (Fig. 5e). A comparison of technical replicates of the same technique yielded a small number of methylation differences, showing that our analysis is picking up relevant differences that are associated with the choice of protocol. (Additional File 2: Fig. S8b). This is an important finding, showing that researchers can obtain different results and be led to different conclusions depending on which WGBS method they use in their study.

Our results highlight that overestimation of mCG values is a common feature of WGBS protocols with amplification, despite their performance in various bias tests. This is explained by our observations that different sources of bias contribute to this effect in the different protocols, and we have summarised our estimates for some of our datasets in Figure 5f.

### Coping with biases and artefacts

We have shown that many of the biases in WGBS datasets reflect themselves in sequence composition of the libraries. To aid with the evaluation of WGBS biases, we have incorporated, as part of the quality control (QC) package of the Bismark program [51], a ‘bam2nuc’ module to quantitate the mono-and di-nucleotide composition in a WGBS dataset against the genomic expected values (Fig. 6a). A depletion of cytosine mono-and dinucleotide content is likely to be a result of BS-induced DNA degradation, whereas cytosine enrichment would indicate poor conversion efficiency; G or AT depletion/enrichment could serve as measures of amplification bias and polymerase preferences. For specific sequences, the publically available GC-content tracks on genome viewers could be used as an indication for likelihood for biases. We have also created and made public (see Declarations: Availability of Data and Materials) strand-specific G and C content wiggle tracks for the mouse genome.

**Figure 6.**
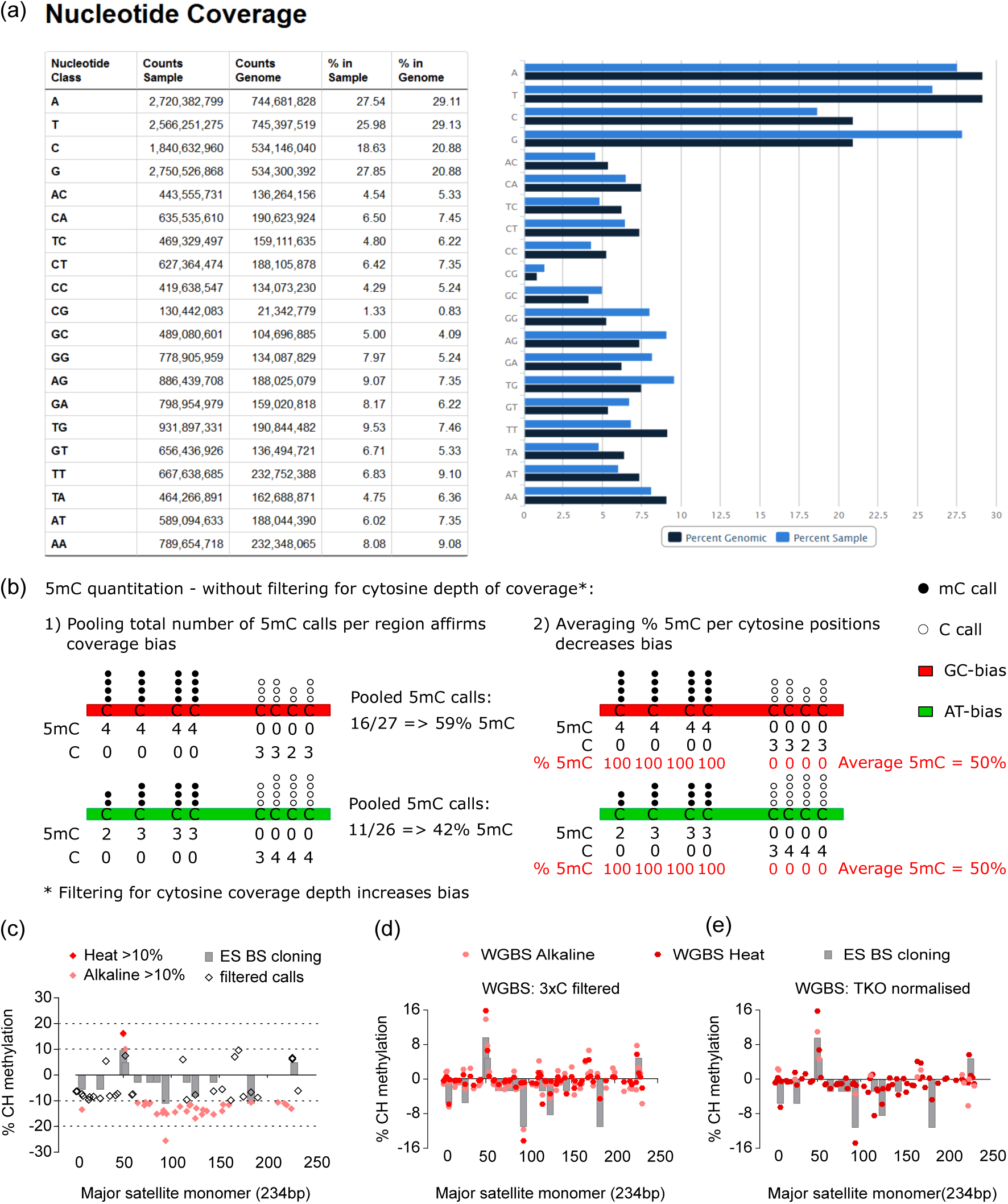
Dealing with biases and artefacts. (**a**) A screenshot fr om the Bismark ‘bam2nuc’ module output, showing base and dinucleotide content in a ‘Heat’ dataset against the genomic expected value. The C base indicates the extent of degradation-caused bias (negative correlation), the G-base and its derivative dinucleotides as well as the A/T bases and dinucleotides show the extent and direction of amplification bias – positive or negative, G(C)-or AT-biasing. (**b**) Comparison of two methylation quantitation strategies to overcome or decrease the effects of GC-or AT-biasing protocols. Counting a total number of methylation calls within a probe (region), irrespective of their position and depth, is compared to calculating methylation values of individual CGs and averaging those for the whole probe. None of those approaches applies initial coverage depth filtering, which is shown in Additional File 2: Figure S9. (**c, d, e**) Non-CG methylation in the major satellite after removing conversion noise by (**c**) setting a 10% or 20% 5mC cut-off threshold value, (**d**) after a bioinformatic filter was applied to remove every read with three or more methylated CH cytosines, and (**e**) after subtracting the background values from an unmethylated genome control (TKO mESC). Positive y-axis values indicate the top strand and negative – the bottom strand.

Another strategy is related to the way methylation is quantitated in regions of interest. Our results showed that polymerase and PCR biases can enrich the sequence output of WGBS datasets either towards GC-rich or towards AT-rich sequences (Fig. 2a). We experimented with common ways to estimate methylation and identified a strategy that is less affected by coverage biases (Fig. 6b). Namely, quantitating methylation for individual cytosines within a region of interest and averaging their values outperforms the alternative practice of pooling all methylation calls within a region. Importantly, the very common practice of selecting for analysis only cytosines with a minimal fold coverage (usually above 5 or 10), results in reinforcement of the coverage bias effects and skews the resulting methylation values (Additional File 2: Fig. S9a).

Incomplete conversion artefacts affect CG methylation and are a subject of over-amplification during PCR, but their most dramatic effect is on non-CG methylation. Spiking of unmethylated DNA of foreign origin during library preparation, such as Lambda or M13 phage DNA, is a useful way to monitor the global conversion efficiency per sample (Additional file 2: Fig. S10a). However, BS conversion resistance is very sequence specific [3,11] (Additional file 2: Fig. 10b) and thus not fully represented by a control of different origin and composition. It is also common to use genomic non-CG context methylation as an indication for conversion efficiency, which may be appropriate for some samples but also risky, given the well documented presence of mCH in mammalian, insect and plant genomes [9,26,27,29,30,41,42,46,52,53]. Different approaches can be applied to distinguish real mCH from artefacts and reduce the weight of false positive methylation calls. Here we compared three strategies to cope with conversion errors, tested on the major satellite consensus and our M13 spike-in controls: 1) removal of CH calls below a threshold methylation value (usually set between 3 and 20 %), 2) filtering of reads with 3 or more consecutive unconverted CH bases (the 3xC filter), and 3) a new approach for normalisation of each cytosine’s methylation against an unmethylated WGBS control of the same genome. Although the first approach is the most commonly used [52,54,55], our results show that it is the least efficient one, since a large number of false positive calls remain above the threshold, which at the same time is likely to remove real methylation calls (Fig. 6c and Additional file 2: Fig. 10b). The second approach is much more efficient in removing the background noise, although a number of conversion resistant GC-rich sites (such as CCWGGs) can pass the filter (Fig. 6d and Additional file 2: Fig. 10c). For the third approach, we used our unmethylated TKO mESC WGBS datasets prepared with ‘Heat’ and ‘Alkaline’ protocols (Additional file 2: Fig. 10d) as background noise controls to subtract from the corresponding WT mES datasets. This substantially reduced the noise from CH context positive calls (shown in Fig. 4e) to levels comparable to those achieved with the classic BS cloning and the 3xC bioinformatic filter (Fig. 6e). The main benefit of this approach is the ability to deal with conversion resistant sequences, such as the ones we observe in the M13 spike-in, which pass through the 3xC filter (Additional file 2: Fig. 10b). Our results demonstrate that WGBS is not a noise-free technique, therefore studies interpreting non-CG methylation should be accompanied with robust controls and have clear strategies for coping with conversion artefacts.

## Discussion

Here we present a comparative analysis between five BS conversion methods and seven WGBS library preparation protocols, dissecting the most common sources of bias. We have evaluated their performance and summarize our results in Table 3.

**Table 3.** Summary of biases affecting the pre-and post-BS WGBS library preparation strategies. For simplicity, pre-BS methods have been divided into GC-biasing (‘Heat’, ‘Alkaline’ and ‘Am-BS’) and AT-biasing (‘KAPA’), and post-BS methods into amplification-free (‘PBAT’) and with amplification (‘ampPBAT’ and ‘EpiGnome’). (+) indicates low bias, (++) medium and (+++) high bias.

| BIAS | Main result figures | PRE-BISULFITE |  | POST-BISULFITE |  |
| --- | --- | --- | --- | --- | --- |
|  |  | GC-biasing | AT-biasing (KAPA) | Amplification-free | With amplification |
| **Degradation bias (C depletion)** | Fig. 1a; Fig. 2a,c | +++ | +++ | + | + |
| **PCR (polymerase) bias** | Fig. 2a-c, Fig. 5f | +++ | + | + | +++ |
| **Modified C bias** | Fig. 1d; Fig. 3d | ++ | ++ | + | ++ |
| **Conversion artefacts** | Fig. 4 | ++ | + | + | + |
| **Global 5mC overestimation** | Fig. 2d | +++ | ++ | + | ++ |

Our findings reveal that WGBS protocols suffer from multiple biases and have a highly variable performance, a fact that has not received due attention to date. The biases lead to overestimated absolute levels of both CG and CH context methylation, skewed relative methylation differences between samples and under- or over-representation of vulnerable genomic regions. These unwanted effects can be modulated by a careful selection of the library preparation strategy and specific conditions during key steps, but are best avoided with an amplification-free approach (Table 3).

Our results show that BS-mediated DNA degradation is the underlying cause for biases in WGBS data. It affects the sequence composition and methylation output through depletion of unmethylated C-rich regions. This effect seems stronger in pre-BS approaches, where DNA is fragmented and adapter-tagged prior to BS conversion. During BS treatment the library undergoes a second fragmentation step, where C-rich unmethylated fragments get excluded from the library pool before they undergo amplification, which introduces a sequence bias. The uneven representation of strands and sequences predisposes increased stochasticity in the first cycles during PCR, resulting in under-or over-representation of certain sequences, irrespective of polymerase GC bias [21], an effect observed in both low and high PCR cycle libraries (Additional File 2: Fig.S3a). Post-BS methods, on the contrary, harness the BS-induced fragmentation by directly starting from high molecular weight DNA to yield the desired fragment size [8,28] and decrease the loss of C-high content. This allows for a more accurate estimation of global methylation levels even after amplification (Fig. 2d), although localised and feature CG methylation values are nevertheless altered after amplification (Fig. 5). Harsher BS conversion conditions such as heat denaturation of DNA and higher BS incubation temperatures (65-70°C) yield a better and more consistent BS conversion efficiency, especially if combined with high molarity of BS (>4 M) and short incubation times (30-90 min) (Fig. 4a and Additional File 2: Fig.10a) [5,8,14]. Longer incubation times have been shown to lead to higher degradation and accumulation of inappropriate conversion (false negatives), without necessarily contributing to conversion efficiency [5,8]. The harsh BS conditions, however, create strong biases with the pre-BS approach when combined with GC-biasing polymerases like the Pfu Turbo Cx, but are the preferred choice with KAPA Uracil+ and the post-BS protocols. The choice of reliable conversion conditions is particularly important for studying non-CG methylation, where the alkaline denaturation, low BS molarity and lower temperatures are likely to yield false positives, which outnumber the real 5mCH signal in a sample (Fig. 4c and 5e).

PCR amplification was found to build on the over-represented methylated sequences and conversion artefacts, thus amplifying on the errors from BS treatment and becoming a major source of bias for both pre-and post-BS methods. The best performance was observed for the amplification-free PBAT approach, where, in addition to the low degradation bias, it showed insignificant CG-context coverage bias and better matched the 5mC levels measured by LC/MS (Fig. 2a-d). The amplification-free output was also least affected by the underlying methylation status (Fig. 3d). Given that a main advantage of PBAT has been its use with very low DNA input [28,29,33], the amplification-free approach should be feasible for most standard applications. Whilst the original PBAT method suffers from lower mapping efficiency due to chimeric reads, these are easily dealt with during the bioinformatic processing [56]. However, it has recently been reported that the original PBAT is affected by different versions of the Illumina HiSeq base calling software, which can affect the estimation of global mCG values [57]. Importantly, an alternative amplification-free WGBS approach called ReBuilT has been published more recently, also showing an improvement in GC-bias [58]. Amplification-free protocols are reported the least biased solution for NGS microbiome analysis, where sequence diversity and (mis)-representation is of high importance, as in WGBS [59].

Classically PCR bias has been associated with enrichment of GC-rich sequences [16–20,23]. The KAPA HiFi family of polymerases have been shown to have low GC-bias and our results show that indeed G, but not C, coverage is improved by using this enzyme (compare ‘Pfu’ and ‘KAPA’ in Fig. 2a-c). Notably, we show that by minimizing amplification artefacts, the overall low bias of KAPA Uracil+ yields global methylation estimates close to those obtained with post-BS approaches (Fig. 2d). Yet at the local level, differences are seen when comparing ‘KAPA’ pre-BS libraries with amplification-free PBAT ones (Fig. 5a-c and Additional File 2: Fig. S6 and S7), which could be due to the preferential degradation of DNA fragments containing unmodified cytosines (Fig. 1d) that are not recovered by pre-BS methods.

A growing number of WGBS datasets are currently available in the public domain and often datasets generated by different labs get used together in one study. Given the presented differences in the methods’ absolute and relative methylation estimates, analysing and comparing data generated by different protocols should be avoided or done with caution in an informed manner, with the biases accounted for during the analysis and interpretation of results. Our new ‘bam2nuc’ module in the QC package of the Bismark software aims to help assess the strength and direction of biases, to avoid the interpretation of purely technical methylation differences as biological. Our data shows that changes up to 20% in DNA methylation can be purely technical.

Our results demonstrate the existence of conversion artefacts ‘noise’ that is particularly relevant in the context of non-CG methylation, especially as the latter becomes increasingly a focus of biological interest [29,30,41,42,46,52,53]. We show that bioinformatic filtering of reads with three or more consecutive unconverted CH cytosines is necessary and more efficient than setting a cut-off threshold value, and should become a standard even in datasets with high overall conversion rates. This tool has now been integrated as an optional ‘filter_non_conversion’ module in the Bismark methylation caller [51]. Alternatively, sequencing a whole genome amplified (WGA) unmethylated sample in addition to the samples of interest can be used to normalise false discovery rates with single base resolution. The latter approach is particularly important for studies of model organisms with very low or questionable methylation levels near the detection limit, such as insects. Such studies, especially reporting methylation in C-rich regions or non-CG context [47], should be backed up by unmethylated genome controls and validated with non-BS methods such as LC/MS [41].

## Conclusions

Our findings establish basic principles for understanding and minimising biases when designing and optimising WGBS strategies. We envisage that, in the current state-of-the-art, the gold standard for WGBS library preparation should evolve towards an amplification-free approach with optimised BS treatment conditions and, where necessary, low-bias DNA polymerases. Such benchmark method would be of great value for the research community and enable researchers from outside fields to always generate methylation data with minimal biases. We hope our results would also encourage the development of newer and better amplification-free protocols. New sequencing technologies, not dependent on BS treatment, will also push the field forward and help obtain degradation bias-free and conversion error-free maps of DNA methylation.

## Methods

### ES cell culture and DNA preparation

The J1 ES cell line (129S4/SvJae) was purchased from ATCC (Cat. SCRC-1010) and the Dnmt1-/-,3a-/-,3b-/-TKO line is a kind gift from Masaki Okano [60]. The TKO was cultured on gelatine without feeders and the J1 on a γ-irradiated pMEF feeder layer, at 37°C and 5% CO_2_ in complete ES medium (DMEM 4500 mg/L glucose, 4 mM L-glutamine and 110 mg/L sodium pyruvate, 15% foetal bovine serum, 100 U of penicillin/100 μg of streptomycin in 100 mL medium, 0.1mM non-essential amino acids, 50 μM β-mercaptoethanol, 10^3^U/ml LIF ESGRO^®^). Mycoplasma tests on cell lines are routinely performed in the lab. Genomic DNA was extracted with DNeasy Blood and Tissue Kit (Qiagen) following manufacturer’s instructions and quantitated via Quant-iT™ PicoGreen dsDNA Assay Kit (Invitrogen). *In vitro* DNA methylation with M.CviPI (New England Biolabs, NEB) was performed on 0.5 – 1.0 μg of TKO mESC genomic DNA, incubated for 2 hours at 37°C, purified with GeneJet PCR Purification kit (Thermo) and quantitated by Quant-iT™ PicoGreen^®^ dsDNA Assay Kit (Invitrogen).

### M13 fragments analysis

M13mp18 (NEB) was used as a template. PCR of C-poor and C-rich fragments (Additional file 2: Table S1) was performed using either a standard dNTP mix (Bioline), or substituting the dCTPs with modified dm5CTPs (10 mM, NEB) or d5hmCTPs (100 mM, Bioline). The PCR was performed with Dream Taq DNA Polymerase (Thermo Scientific) (50 μl volume, 200 nM primer, 200 μM dNTPs, 1.25 units enzyme) with an initial step at 95°C for 2 minutes, 35 cycles of 30 seconds at 95°C, 20 seconds at 57°C, and 30 seconds (C, 5hmC) or 5 minutes (5mC) at 72°C, with a 7 minute final step at 72°C (primers are listed in Additional file 2: Table S2). All PCR products were verified on a DNA resolving 2% agarose gel, purified with GeneJet PCR Purification kit (Thermo Scientific) and quantitated by both Quant-iT™ PicoGreen dsDNA Assay Kit (Invitrogen) and Agilent 2100 Bioanalyzer. Identical aliquots were prepared from each fragment for BS treatment with the different protocols and an aliquot each was kept as an input control. BS treated fragments were eluted in the same final volume as the input and quantitated for recovery 3 to 4 times for each sample. Each BS treatment was repeated twice.

### Bisulfite conversion of DNA

Genomic DNA and purified M13-derived fragments were treated with sodium bisulfite using all of the following kits: EpiTect Bisulfite kit from Qiagen (FFTP protocol), Imprint DNA Modification kit from Sigma-Aldrich (1-step and 2-step) and EZ DNA Methylation Kit (Zymo Research) according to manufacturer’s instructions. The *in vitro* M.CviPI-methylated TKO DNA was converted with the EpiTect Bisulfite kit. A minor modification was applied for all samples treated with the EpiTect kit: the 5 hour incubation programme was run twice (10 hours) following a commonly accepted practice [26,53,61,62]. Conversion with 9 M ammonium bisulfite was performed at 70°C for 30 minutes as in Hayatsu et al. [25]; 50% Ammonium Hydrogen Sulfite Solution was purchased from Wako Chemicals GmbH, the rest of the reagents were supplied by Sigma-Aldrich. Half of the samples converted with the 1-step (‘heat’) and 2-step (‘alkaline’) Imprint DNA Modification kit and the 9M ammonium bisulfite procedure were purified with Amicon Ultra 0.5mL Ultracel 30k filters (Millipore), with the clean-up reagents and following the manufacturer’s purification instructions of the TrueMethyl kit v1 (Cambridge Epigenetix).

### Purification, cloning and BS sequencing of the major satellite repeat

J1 (WT) and TKO ES [60] genomic DNA was bisulfite converted using EpiTect Bisulfite Kit (Qiagen) as explained previously. The major satellite was amplified with HotStart Taq (Qiagen) in a mixture of 200 nM primer, 200 μM dNTPs, 2 mM MgCl_2_, 1.0 unit of enzyme at 94°C for 15 minutes, 35 cycles of 20 seconds at 94°C, 20 seconds at 55°C, and 20 seconds at 72°C, and a final step at 72°C for 3 minutes. DNA fragments spanning over one repeat (370bp) were excised from 2% agarose gels and purified with a MinElute Gel Extraction kit (Qiagen) following the kit protocol. The fragments were cloned into pGEM-T using the pGEM-T Easy Vector Kit (Promega) and transformed into Invitrogen’s Subcloning Efficiency DH5α Competent Cells according to manufacturer’s instructions. Positive clones were selected on LB plates containing 100 μg/ml ampicillin and covered with X-gal (40 mg/ml). Colonies were screened with Roche’s Taq DNA Polymerase (25 μl volume, 300 nM primer, 200 μM dNTPs, 1.25 units enzyme) at 94°C for 10 minutes, 35 cycles of 30 seconds at 94°C, 30 seconds at 55°C, and 30 seconds at 72°C, with a final 72°C for 10 minutes and sent for Sanger sequencing at Beckman Coulter Genomics. All oligonucleotides are listed in Additional file 2: Table S2. Methylation of the sequenced clones was analysed with QUMA [63] and plotted with a custom made R script.

### Mass spectrometry

Untreated or BS treated genomic DNA (0.3-1 μg) was digested with a DNA Degradase Plus™ (Zymo Research) for 3 hours at 37°C according to manufacturer’s instructions. Approximately 50-100 pg per sample were analysed by LC-MS/MS on a Thermo Q-Exactive mass spectrometer coupled to a Proxeon nanoLC. Three replicates of each sample were analysed and the amounts of C, 5mC, 5hmC and U and T were quantified relative to external standards. Recovery of BS treated genomic DNA for the different BS conversion protocols and clean-up procedures was assessed in the same way as for the M13 fragments, but quantitated with the LC/MS.

### Library preparation and next generation sequencing

Approximately 250 ng genomic DNA was fragmented via sonication with a Covaris E220 instrument with the 300 bp programme, and spiked in 1:10,000 with a 2 kb unmethylated PCR fragment from M13mp18 (New England Biolabs). Early Access Methylation Adaptor Oligos (Illumina) were ligated to the fragmented DNA with the NEB Next DNA Library Prep Master Mix Set for Illumina (E6040), according to the manufacturer’s instructions and purified after each step with Agencourt^®^ AMPure^®^ XP beads. Bisulfite treatment was performed as described above in ‘Bisulfite conversion of DNA’, using all listed methods, except for the ammonium bisulfite protocol. The BS converted libraries were amplified using PfuTurbo Cx Hotstart DNA Polymerase (Agilent Technologies): 300 μM dNTPs, 400 nM indexed adaptor-specific primers[19], 2.5 units enzyme, with an initial step at 98°C for 30 seconds, 15 cycles of 98°C for 10 seconds, 65°C for 30 seconds, and 72°C for 30 seconds, with a final elongation step at 72°C for 5 minutes. Library quality control was performed with an Agilent 2100 Bioanalyzer and quantity determined via KAPA Library Quantification Kit (KAPA Biosystems). For the unmethylated DNA controls, library preparation was performed in the same way, with the following modifications: 1.0 μg of DNA was sonicated and adaptor-ligated with Illumina TruSeq indexed adaptors, no BS conversion was performed, and amplification was done with the NEB Next 2x Phusion mix for 6 cycles, following manufacturer’s instructions.

Paired-end 100 bp NGS was performed on an Illumina HiSeq 2000 system at the Bespoke Facility at the Wellcome Trust Sanger Institute.

### WGBS data mapping and quality analysis

Data from both BS converted and non-BS converted datasets, was trimmed with Trim Galore and raw data quality analysis performed with FastQC [64]. Mapping was carried out with Bismark [51] to NCBIM37 and GRCm38 builds for the mouse genome, GRCh38 for the human genome, HS3.3 for the ant *Harpegnathos saltator*, GCA_000297895.1 for the Pacific oyster *Crassostrea gigas*, Chinese hamster reference sequence in Ensembl for the CHO cells, and assembly scaffolds for the ant *Dinoponera quadriceps*. For consistency and to reduce error, the non BS converted datasets were also mapped with Bismark. All alignments were performed with high stringency allowing for only one base mismatch (n=1) and mapped data was deduplicated before analyses. For PBAT libraries, all mapping errors from chimeric reads and M-bias were taken into consideration upon processing and first 4 bases from each read were excluded from the analysis [56]. All datasets were deduplicated, consistent with common analyses pipelines, although this decreased the sequence and methylation bias, normally stronger in the raw duplicated data (unpublished observation). Some of the analyses, however, such as telomere and major satellite, were performed on raw (duplicated) data, since those sequences are not mappable.

### Read coverage analysis

After processing with Bismark, read coverage depth was analysed with SeqMonk [65]. Custom genomes were created for mtDNA, satellite repeat and M13, and the reads were aligned preserving the strand information. Because of its repetitive nature, the data for major satellite was not deduplciated following alignment. Read coverage was assessed from the total read count for forward or reverse strands in each custom-built genome.

For the mouse CGI coverage, regions with unusually high read coverage (1kb genomic windows with more than 1000 reads), likely to represent alignment artefacts were excluded from the analysis. The CGI coverage analysis was performed in SeqMonk via a relative trend plot over CGIs (coordinates from Illingworth et al. 2010 [66]), including all reads, ‘forced to be relative’ option chosen and allowing for 1kb flanks.

### Methylation analysis of WGBS

CG methylation quantitation was performed in SeqMonk with the integrated bisulfite analysis pipeline. Regions likely to attract alignment artefacts (having more than 1000 reads over 1kb genomic windows) were excluded from the analyses. CG methylation analysis on genic and regulatory features was quantitated on the individual replicate datasets via probes created over each feature, without setting cytosine coverage thresholds but requiring minimum 3 observations in order to include a probe. This analysis was undertaken only for mouse ESC datasets, for which public datasets were available for all compared protocols, except for Am-BS (see Additional file 1 for details). Promoters were defined as -1000 + 200bp from the TSS, DMR coordinates were obtained from Seisenberger et al. [40], CGI coordinates from Illingworth et al. 2010 [66], active, poised and primed enhancers from Creyghton et al. 2010 [67], super-enhancers from Whyte et al. 2013 [68], ES-specific LMRs from Stadler et al. 2011 [50], Yamanaka factor binding sites from Chen et al. 2008 [69] and transcription factor binding sites (TFBS) from UCSC (Caltech annotation). Exons and introns were defined with Ensembl-derived coordinates integrated in SeqMonk. IAP, LINE, LTR, Satellite and SINE coordinates were derived from UCSC [70].

The genome-wide analysis (scatter plots) was performed for two groups of datasets: 1) the panel of published mouse ESC datasets from six WGBS methods generated by different studies or labs and used for the feature analyses (see Additional file 1), and 2) datasets generated by the same lab with PBAT and Heat BS-seq from four different biological samples (mESC, sperm, blastocyst and oocyte) [29]. The first group provides comparison between all protocols (except for Am-BS due to unavailability), while the second set serves to validate the observations from the first group with the difference that it should not be affected by potential sample strain or batch differences. For the first group probes were made over non-overlapping 50-cytosine containing tiles (i.e. measurement windows) over the PBAT mESC datasets and quantitated for all remaining datasets as pooled replicates. For the second set of datasets 150C-containing measurement windows were made over each corresponding PBAT dataset and quantitated over pooled or individual replicates for the PBAT vs Heat BS-seq comparison or for the inter-replicate comparison, respectively.

The relative methylation analysis was performed with PBAT and Heat BS-seq datasets from Kobayashi et al. [29], used for the whole genome analysis above. We made probes over 150-cytosine containing tiles over the whole genome for the mESC PBAT dataset (due to lowest coverage) and quantitated CG methylation for these probes in PBAT sperm, as well as in the BS-seq datasets for both samples. We then selected regions with 20% CG methylation difference between sperm and ESC in both the PBAT and BS-seq datasets and plotted the regions obtained with PBAT onto the BS-seq plot and vice versa. Overlapping differentially methylation probes between the PBAT and BS-seq lists were quantitated in SeqMonk and plotted as venn diagrams with R.

CH filtering of major satellite and M13 was done by removing every read containing more than three unconverted CH cytosines. The script can be found in the provided Github deposition and also integrated as ‘filter_non_conversion’ module in the Bismark package v0.17.0 [51]. All analyzed datasets and the number of replicates per protocol are listed in Additional file 1.

### Sequence composition analyses

All described sequence composition analyses were performed with SeqMonk and custom-made Perl and R scripts. All scripts can be accessed from the Github deposition directory provided under Declarations.

Dinucleotide coverage was assessed through the quantitation of the total number of dinucleotide instances within the mapped data of each dataset. It was plotted against the expected occurrence of each dinucleotide derived from the relevant annotated genome (see Additional file 3 for genomic references). This analysis is integrated as ‘bam2nuc’ module in Bismark v0.16.0 [51].

For the telomere analyses we used only raw data reads, since the tandem hexamer units of the telomere are not mappable. We quantitated the number of occurrences of each hexamer (or heptamer for *A. thaliana*) per read, as follows: TTAGGG (G-strand hexamer) or TTTTAA (BS converted version of the CCCTAA C-strand hexamer) for BS-seq and EpiGnome datasets; the reversed sequences CCCTAA(A) or (T)TTAAAA for PBAT and ampPBAT datasets (heptamers, in parentheses, were used for *A. thaliana*); TTAGGG, CCCTAA or TTTTAA for the non-BS converted control. The TTTTAA hexamer was quantitated in the non-BS converted control in order to assess the original genomic occurrence of this non-telomere sequence prior to BS conversion. Read lengths varied between 30-100bp for BS-seq datasets, 44bp, 76bp or 121bp for PBAT and 100bp for the non-BS control, hence the variation in number of units in Fig. 1c and S1b. Our quantitation revealed that TTTTAA and (T)TTAAAA occurred mostly in one to five units per read before BS conversion, indicating that the majority of those reads are native to the genome and do not derive from the telomere repeat. The telomere hexamers (T)TTAGGG and CCCTAA(A), however, were present in higher numbers per read. In order to exclude the non-telomere derived TTTTAA and (T)TTAAAA hexa/hepta-mers from the BS treated data, therefore, all reads containing less than 5 units per read we removed from the analyses.

For the correlation of read coverage with C and G content, we generated non-overlapping 100bp running window tiles over the whole mouse genome. Regions likely to attract alignment artefacts (defined previously as having more than 1000 reads over 1kb genomic windows) were excluded from the analysis. Read count of each 100bp-tile was quantitated and base composition was extracted from the genomic sequence and not the actual read sequence, where cytosines are BS converted to thymines. The tiles were grouped in 100 bins by their G or C content and the mean read count per tile, normalised to the total read count per dataset, was plotted against the % G or C.

### Statistics and sample size

Statistical analyses were performed with GraphPad Prism 6.0. Error bars represent standard deviation (s.d.) or standard error of the mean (s.e.m), as described in each figure legend, and indicate whether biological or technical replicates are compared, respectively. Where possible, exact p-values are stated in the figures or figure legends, otherwise star-symbols are used with the corresponding p-value ranges indicated. Where applicable, value distribution was tested with D’Agostino-Pearson normality test. All LC/MS analyses were performed on two separate BS conversion experiments for each method, with 3 replicate samples each (i.e. 6 in total), and a minimum of three LC/MS measurements per sample. For WGBS, we performed our analyses on a panel of datasets either sequenced for this study (2 to 3 replicates each) or publically available, generated for different studies or by different laboratories (see details in Additional file 1). Where possible, information on number of samples is provided in the figures or figure legends.

### List of abbreviations

5mC: 5-methylcytosine; 5hmC: 5-hydroxymethylcytosine; BS: bisulfite; WGBS: whole genome bisulfite sequencing; RRBS: reduced representation bisulfite sequencing; DNA: deoxy-ribonucleic acid; NGS: next generation sequencing; PCR: polymerase chain reaction; PBAT: post-bisulfite adaptor-tagging; C: cytosine; LC/MS: liquid chromatography with mass-spectrometry; mESC: mouse embryonic stem cells; TMAC: tetramethylammonium chloride; SNP: single nucleotide polymorphism; indel: insertion and deletion-caused mutations; s.d.: standard deviation; s.e.m.: standard error of the mean; TKO: triple-Dnmt knock-out cell line; CGI: CG-island; (i)DMR: (imprinted) differentially methylated region; QC: quality control; WGA: whole genome amplification.

## Declarations

### Ethics approval and consent to participate

Not applicable

### Consent for publication

Not applicable

### Availability of data and materials

All datasets generated in this study are listed in Additional file 1 and deposited in NCBI’s GEO under the accession number <u>GSE77961</u> [45]. Modules ‘bam2nuc’ and ‘filter_non_conversion’ are part of the Bismark suite [51], which is released under the GNU GPL v3 license. All Perl and R scripts, including those for generating mouse genome C-and G-content wiggle tracks, are deposited in GitHub [71].

### Author’s contributions

N.O., M.R.B. and W.R. conceived the study and designed experiments. N.O. performed experiments and data interpretation, N.O., F.K. and S.A. designed and performed the bioinformatic analyses, D.O. performed mass spectrometry, R.V.B. provided unpublished data, N.O., M.R.B. and W.R. wrote the manuscript. All authors read and approved the final manuscript.

## Acknowledgements

We thank Anne Segonds-Pichon for help with statistical analyses, Wendy Dean and all Reik lab members for helpful discussions, Judith Webster for assistance with mass spectrometry and the Bespoke Facility at the Wellcome Trust Sanger Institute for Illumina sequencing.

## Additional files

Additional file 1: an Excel spreadsheet with three tabs listing 1) datasets generated in this study, 2) datasets used from the publically domain, together with key parameters, and 3) number of datasets used per analysis.

Additional file 2: a PDF file with supplementary Tables S1-3 and supplementary Figures S1-10.

Additional file 3: an Excel spreadsheet with reference genome compositions.

## References

1 Frommer M, Mcdonald LE, Millar DS, Collist CM, Wattt F, Griggt GW, et al. A genomic sequencing protocol that yields a positive display of 5-methylcytosine residues in individual DNA strands. PNAS. 1992;89:1827–31.

2 Grunau C, Clark SJ, Rosenthal a. Bisulfite genomic sequencing: systematic investigation of critical experimental parameters. Nucleic Acids Res. 2001;29:E65–5.

3 Warnecke PM, Stirzaker C, Song J, Grunau C, Melki JR, Clark SJ. Identification and resolution of artifacts in bisulfite sequencing. Methods. 2002;27:101–7.

4 Harris RA, Wang T, Coarfa C, Nagarajan RP, Hong C, Downey SL, et al. Comparison of sequencing-based methods to profile DNA methylation and identification of monoallelic epigenetic modifications. Nat. Biotechnol. Nature Publishing Group; 2010;28:1097–105.

5 Genereux DP, Johnson WC, Burden AF, Stöger R, Laird CD. Errors in the bisulfite conversion of DNA: modulating inappropriate- and failed-conversion frequencies. Nucleic Acids Res. 2008;36:e150.

6 Warnecke PM, Stirzaker C, Melki JR, Millar DS, Paul CL, Clark SJ. Detection and measurement of PCR bias in quantitative methylation analysis of bisulphite-treated DNA. Nucleic Acids Res. 1997;25:4422–6.

7 Raizis AM, Schmitt F, Jost JP. A bisulfite method of 5-methylcytosine mapping that minimizes template degradation. Anal. Biochem. 1995. p. 161–6.

8 Holmes EE, Jung M, Meller S, Leisse A, Sailer V, Zech J, et al. Performance Evaluation of Kits for Bisulfite-Conversion of DNA from Tissues, Cell Lines, FFPE Tissues, Aspirates, Lavages, Effusions, Plasma, Serum, and Urine. PLoS One. 2014;9:1–15.

9 Henderson IR, Chan SR, Cao X, Johnson L, Jacobsen SE. Accurate sodium bisulfite sequencing in plants. Epigenetics. 2010;5:47–9.

10 Hayatsu H, Tsuji K, Negishi K. Does urea promote the bisulfite-mediated deamination of cytosine in DNA? Investigation aiming at speeding-up the procedure for DNA methylation analysis. Nucleic Acids Symp. Ser. (Oxf). 2006;69–70.

11 Harrison J, Stirzaker C, Clark SJ. Cytosines adjacent to methylated CpG sites can be partially resistant to conversion in genomic bisulfite sequencing leading to methylation artifacts. Anal. Biochem. 1998;264:129–32.

12 Chhibber A, Schroeder BG. Single-molecule polymerase chain reaction reduces bias: application to DNA methylation analysis by bisulfite sequencing. Anal. Biochem. 2008;377:46–54.

13 Voss KO, Roos KP, Nonay RL, Dovichi NJ. Combating PCR Bias in Bisulfite-Based Cytosine Methylation Analysis. Betaine-Modified Cytosine Deamination PCR. Anal. Chem. 1998;70:3818–23.

14 Shiraishi M, Hayatsu H. High-speed conversion of cytosine to uracil in bisulfite genomic sequencing analysis of DNA methylation. DNA Res. 2004;11:409–15.

15 Candiloro ILM, Mikeska T, Dobrovic A. Assessing alternative base substitutions at primer CpG sites to optimise unbiased PCR amplification of methylated sequences. Clin. Epigenetics. Clinical Epigenetics; 2017;9:1–9.

16 Aird D, Ross MG, Chen W-S, Danielsson M, Fennell T, Russ C, et al. Analyzing and minimizing PCR amplification bias in Illumina sequencing libraries. Genome Biol. BioMed Central Ltd; 2011;12:R18.

17 Dabney J, Meyer M. Length and GC-biases during sequencing library amplification: A comparison of various polymerase-buffer systems with ancient and modern DNA sequencing libraries. Biotechniques. 2012;52.

18 Ross MG, Russ C, Costello M, Hollinger A, Lennon NJ, Hegarty R, et al. Characterizing and measuring bias in sequence data. Genome Biol. 2013;14:R51.

19 Quail MA, Otto TD, Gu Y, Harris SR, Skelly TF, McQuillan JA, et al. Optimal enzymes for amplifying sequencing libraries. Nat. Methods. 2011;9:10–1.

20 Oyola SO, Otto TD, Gu Y, Maslen G, Manske M, Campino S, et al. Optimizing Illumina next-generation sequencing library preparation for extremely AT-biased genomes. BMC Genomics. BioMed Central Ltd; 2012;13:1.

21 Kebschull JM, Zador AM. Sources of PCR-induced distortions in high-throughput sequencing data sets. Nucleic Acids Res. 2015;43:e143–e143.

22 Kozarewa I, Ning Z, Quail M a, Sanders MJ, Berriman M, Turner DJ, et al. Amplification-free Illumina sequencing-library preparation facilitates improved mapping and assembly of (G+C)-biased genomes. Nat. Methods. 2009;6:291–5.

23 Ji L, Sasaki T, Sun X, Ma P, Lewis ZA, Schmitz RJ. Methylated DNA is over-represented in whole-genome bisulfite sequencing data. Front. Genet. 2014;5:1–10.

24 Tanaka K, Okamoto A. Degradation of DNA by bisulfite treatment. Bioorg. Med. Chem. Lett. 2007;17:1912–5.

25 Hayatsu H, Negishi K, Shiraishi M. DNA methylation analysis: speedup of bisulfite-mediated deamination of cytosine in the genomic sequencing procedure. Proc. Japan Acad. Ser. B. 2004;80:189–94.

26 Lister R, O’Malley RC, Tonti-Filippini J, Gregory BD, Berry CC, Millar AH, et al. Highly integrated single-base resolution maps of the epigenome in Arabidopsis. Cell. 2008;133:523–36.

27 Cokus SJ, Feng S, Zhang X, Chen Z, Merriman B, Haudenschild CD, et al. Shotgun bisulphite sequencing of the Arabidopsis genome reveals DNA methylation patterning. Nature. 2008;452:215–9.

28 Miura F, Enomoto Y, Dairiki R, Ito T. Amplification-free whole-genome bisulfite sequencing by post-bisulfite adaptor tagging. Nucleic Acids Res. 2012;40:e136.

29 Kobayashi H, Sakurai T, Imai M, Takahashi N, Fukuda A, Yayoi O, et al. Contribution of intragenic DNA methylation in mouse gametic DNA methylomes to establish oocyte-specific heritable marks. PLoS Genet. 2012;8:e1002440.

30 Shirane K, Toh H, Kobayashi H, Miura F, Chiba H, Ito T, et al. Mouse Oocyte Methylomes at Base Resolution Reveal Genome-Wide Accumulation of Non-CpG Methylation and Role of DNA Methyltransferases. PLoS Genet. 2013;9:e1003439.

31 Smallwood SA, Lee HJ, Angermueller C, Krueger F, Saadeh H, Peat J, et al. Single-cell genome-wide bisulfite sequencing for assessing epigenetic heterogeneity. Nat. Methods. 2014;11:817–20.

32 Peat JR, Dean W, Clark SJ, Krueger F, Smallwood SA, Ficz G, et al. Genome-wide Bisulfite Sequencing in Zygotes Identifies Demethylation Targets and Maps the Contribution of TET3 Oxidation. Cell Rep. 2014;9:1990–2000.

33 Okae H, Chiba H, Hiura H, Hamada H, Sato A, Utsunomiya T, et al. Genome-Wide Analysis of DNA Methylation Dynamics during Early Human Development. PLoS Genet. 2014;10:1–12.

34 Dai H-Q, Wang B-A, Yang L, Chen J-J, Zhu G-C, Sun M-L, et al. TET-mediated DNA demethylation controls gastrulation by regulating Lefty–Nodal signalling. Nature. Nature Publishing Group; 2016;538:528–32.

35 Khanna A, Czyz A, Syed F. EpiGnome TM Methyl-Seq Kit: a novel post – bisulfite conversion library prep method for methylation analysis. Nat. Publ. Gr. Nature Publishing Group; 2013;10:iii–iv.

36 Farlik M, Sheffield NC, Nuzzo A, Datlinger P, Schönegger A, Klughammer J, et al. Single-Cell DNA Methylome Sequencing and Bioinformatic Inference of Epigenomic Cell-State Dynamics. Cell Rep. 2015;10:1386–97.

37 Ficz G, Branco MR, Seisenberger S, Santos F, Krueger F, Hore T a, et al. Dynamic regulation of 5-hydroxymethylcytosine in mouse ES cells and during differentiation. Nature. 2011;473:398–402.

38 Feichtinger J, Hernandez I, Fischer C, Hanscho M, Auer N, Hackl M, et al. Comprehensive genome and epigenome characterization of CHO cells in response to evolutionary pressures and over time. Biotechnol. Bioeng. 2016;113:2241–53.

39 Milagre I, Stubbs TM, King MR, Spindel J, Santos F, Krueger F, et al. Gender Differences in Global but Not Targeted Demethylation in iPSC Reprogramming. Cell Rep. 2017;18:1079–89.

40 Seisenberger S, Andrews S, Krueger F, Arand J, Walter J, Santos F, et al. The dynamics of genome-wide DNA methylation reprogramming in mouse primordial germ cells. Mol. Cell. 2012;48:849–62.

41 Patalano S, Vlasova A, Wyatt C, Ewels P, Camara F, Ferreira PG, et al. Molecular signatures of plastic phenotypes in two eusocial insect species with simple societies. PNAS. 2015;112:13970–13975.

42 Bonasio R, Li Q, Lian J, Mutti NS, Jin L, Zhao H, et al. Genome-wide and caste-specific DNA methylomes of the ants camponotus floridanus and harpegnathos saltator. Curr. Biol. 2012;22:1755–64.

43 Wang X, Li Q, Lian J, Li L, Jin L, Cai H, et al. Genome-wide and single-base resolution DNA methylomes of the Pacific oyster Crassostrea gigas provide insight into the evolution of invertebrate CpG methylation. BMC Genomics. 2014;15:1119.

44 Raine A, Manlig E, Wahlberg P, Syvanen A-C, Nordlund J. SPlinted Ligation Adapter Tagging (SPLAT), a novel library preparation method for whole genome bisulphite sequencing. Nucleic Acids Res. 2017;45.

45 Olova N, Krueger F, Andrews SR, Branco MR, Reik W. Comparison of whole-genome bisulfite sequencing library preparation strategies identifies sources of biases affecting DNA methylation data [Internet]. GSE77961. NCBI GEO; 2017. Available from: https://www.ncbi.nlm.nih.gov/geo/query/acc.cgi?acc=GSE77961

46 Arand J, Spieler D, Karius T, Branco MR, Meilinger D, Meissner A, et al. In vivo control of CpG and non-CpG DNA methylation by DNA methyltransferases. PLoS Genet. 2012;8:e1002750.

47 Takayama S, Dhahbi J, Roberts A, Mao G, Heo SJ, Pachter L, et al. Genome methylation in D. melanogaster is found at specific short motifs and is independent of DNMT2 activity. Genome Res. 2014;24:821–30.

48 Ficz G, Hore TA, Santos F, Lee HJ, Dean W, Arand J, et al. FGF Signaling Inhibition in ESCs Drives Rapid Genome-wide Demethylation to the Epigenetic Ground State of Pluripotency. Cell Stem Cell. 2013;13:351–9.

49 Berrens R V., Andrews S, Spensberger D, Santos F, Dean W, Gould P, et al. An endosiRNA-Based Repression Mechanism Counteracts Transposon Activation during Global DNA Demethylation in Embryonic Stem Cells. Cell Stem Cell. 2017;21:694–703.e7.

50 Stadler MB, Murr R, Burger L, Ivanek R, Lienert F, Schöler A, et al. DNA-binding factors shape the mouse methylome at distal regulatory regions. Nature. 2011;480:490–5.

51 Krueger F, Andrews SR. Bismark: a flexible aligner and methylation caller for Bisulfite-Seq applications. Bioinformatics. 2011;27:1571–2.

52 Guo W, Chung W-Y, Qian M, Pellegrini M, Zhang MQ. Characterizing the strand-specific distribution of non-CpG methylation in human pluripotent cells. Nucleic Acids Res. 2013;42:1–8.

53 Laurent L, Wong E, Li G, Hodges E, Smith AD, Kendall J, et al. Dynamic changes in the human methylome during differentiation. Genome Res. 2010;320–31.

54 Raddatz G, Guzzardo PM, Olova N, Fantappié MR, Rampp M, Schaefer M, et al. Dnmt2-dependent methylomes lack defined DNA methylation patterns. Proc. Natl. Acad. Sci. U. S. A. 2013;110:8627–31.

55 Xie W, Barr CL, Kim A, Yue F, Lee AY, Eubanks J, et al. Base-resolution analyses of sequence and parent-of-origin dependent DNA methylation in the mouse genome. Cell. Elsevier Inc.; 2012;148:816–31.

56 Krueger F. PBAT libraries may generate chimaeric read pairs [Internet]. QC Fail. 2016. Available from: https://sequencing.qcfail.com/articles/pbat-libraries-may-generate-chimaeric-read-pairs/

57 Toh H, Shirane K, Miura F, Kubo N, Ichiyanagi K, Hayashi K, et al. Software updates in the Illumina HiSeq platform affect whole-genome bisulfite sequencing. BMC Genomics. BMC Genomics; 2017;18:31.

58 McInroy GR, Beraldi D, Raiber E-A, Modrzynska K, van Delft P, Billker O, et al. Enhanced Methylation Analysis by Recovery of Unsequenceable Fragments. PLoS One. 2016;11:e0152322.

59 Jones MB, Highlander SK, Anderson EL, Li W, Dayrit M, Klitgord N, et al. Library preparation methodology can influence genomic and functional predictions in human microbiome research. Proc. Natl. Acad. Sci. U. S. A. 2015;

60 Tsumura A, Hayakawa T, Kumaki Y, Takebayashi S, Sakaue M, Matsuoka C, et al. Maintenance of self-renewal ability of mouse embryonic stem cells in the absence of DNA methyltransferases Dnmt1, dnmt3a and Dnmt3b. Genes to Cells. 2006;11:805–14.

61 Bock C, Tomazou EM, Brinkman AB, Müller F, Simmer F, Gu H, et al. Quantitative comparison of genome-wide DNA methylation mapping technologies. Nat. Biotechnol. Nature Publishing Group; 2010;28:1106–14.

62 Meissner A, Mikkelsen TS, Gu H, Wernig M, Hanna J, Sivachenko A, et al. Genome-scale DNA methylation maps of pluripotent and differentiated cells. Nature. 2008;454:766–70.

63 Kumaki Y, Oda M, Okano M. QUMA: quantification tool for methylation analysis. Nucleic Acids Res. 2008;36:170–5.

64 Andrews S. Babraham Institute Bioinformatics. FastQC. [Internet]. 2010. Available from: http://www.bioinformatics.babraham.ac.uk/projects/fastqc/

65 Andrews S. Babraham Institute Bioinformatics. Seqmonk. [Internet]. 2007. Available from: http://www.bioinformatics.babraham.ac.uk/projects/seqmonk/

66 Illingworth RS, Gruenewald-Schneider U, Webb S, Kerr ARW, James KD, Turner DJ, et al. Orphan CpG islands identify numerous conserved promoters in the mammalian genome. PLoS Genet. 2010;6:e1001134.

67 Creyghton MP, Cheng AW, Welstead GG, Kooistra T, Carey BW, Steine EJ, et al. Histone H3K27ac separates active from poised enhancers and predicts developmental state. Proc. Natl. Acad. Sci. U. S. A. 2010;107:21931–6.

68 Whyte W a, Orlando D a, Hnisz D, Abraham BJ, Lin CY, Kagey MH, et al. Master transcription factors and mediator establish super-enhancers at key cell identity genes. Cell. Elsevier Inc.; 2013;153:307–19.

69 Chen X, Xu H, Yuan P, Fang F, Huss M, Vega VB, et al. Integration of External Signaling Pathways with the Core Transcriptional Network in Embryonic Stem Cells. Cell. 2008;133:1106–17.

70 Rosenbloom KR, Armstrong J, Barber GP, Casper J, Clawson H, Diekhans M, et al. The UCSC Genome Browser database: 2015 update. Nucleic Acids Res. 2015;43:D670–81.

71 Krueger F, Andrews SR, Olova N. BS_bias scripts [Internet]. GitHub. 2016. Available from: https://github.com/NellyOlova/BS_biass

